# Maternal obesity accelerates cardiomyocyte maturation and programs adult heart dysfunction in offspring

**DOI:** 10.64898/2026.10.07.757474

**Authors:** Dounia Farhi, Gilles Dilanian, Erminia Donnarumma, Nicolas Pataluch, Louis Montagne, Camille Blandin, Nathalie Mougenot, Timothy Wai, Céline Galés, Elise Balse

## Abstract

**Rationale:** Maternal obesity increases cardiovascular risk in offspring, but the developmental mechanisms linking early-life metabolic stress to adult heart dysfunction remain poorly understood, and whether maternal obesity alters the postnatal maturation of the cardiomyocyte itself has not been addressed.

**Objective:** To determine whether maternal high-fat diet alters the developmental trajectory of postnatal cardiomyocyte maturation and its functional consequences in adult offspring.

**Methods:** In a rat model of maternal high-fat diet, longitudinal echocardiographic and invasive hemodynamic phenotyping were combined with transcriptomic, lipidomic, and ultrastructural analyses across four postnatal stages.

**Results:** Offspring exposed to maternal high-fat diet displayed accelerated postnatal cardiac maturation, with premature activation of contractile, calcium-handling and metabolic gene programs, concentric hypertrophy, and enhanced systolic performance in juvenile life. This precocious maturation was not sustained: by adulthood, ejection fraction and intrinsic myocardial contractility progressively declined, coinciding with disorganization of sarcomeres, intercalated discs, and excitation-contraction coupling proteins, in the absence of fibrosis, lipid accumulation or major mitochondrial respiratory dysfunction.

**Conclusions:** These findings reveal accelerated cardiomyocyte maturation as a developmental mechanism linking maternal obesity to long-term cardiac vulnerability, with cardiomyocyte cytoarchitecture disruption emerging as an early hallmark that precedes overt systolic failure.

## INTRODUCTION

The prevalence of obesity among women of reproductive age has risen dramatically over the past decades, making maternal obesity one of the most pressing public health challenges worldwide^1^. Beyond increasing the risk of gestational complications, maternal obesity has profound consequences for the long-term health of the offspring^2^. Children born to mothers with obesity exhibit an increased risk of obesity, type 2 diabetes and cardiovascular disease later in life^3^. This concept is consistent with the Developmental Origins of Health and Disease (DOHaD) hypothesis, which proposes that environmental perturbations during early development have lifelong consequences for organ function and disease susceptibility^4^. Despite compelling epidemiological and experimental evidence, the molecular mechanisms linking maternal obesity to cardiovascular disease in the offspring remain largely unknown.

According to the DOHaD paradigm, environmental insults occurring during critical developmental windows can permanently modify organ development and increase disease susceptibility in adulthood^5,6^. The heart undergoes one of its most dynamic periods of maturation during late fetal and early postnatal life. During this period, cardiomyocytes irreversibly withdraw from the cell cycle, undergo hypertrophic growth, assemble highly organized sarcomeres, establish mature intercalated discs and progressively remodel their metabolism toward oxidative fatty acid utilization^7,8^. This maturation program extends into a late postnatal step, during which the cardiomyocyte lateral membrane acquires its mature three-dimensional surface-crest organization together with subsarcolemmal mitochondrial clustering^9,10^. Because these events are tightly coordinated, even subtle alterations in their timing may have profound and lasting consequences for cardiac function.

A growing body of evidence indicates that maternal obesity alters cardiac development before birth and predisposes offspring to cardiac dysfunction later in life. Animal studies have reported fetal cardiac hypertrophy^11,12^, altered myocardial metabolism and impaired ventricular function following maternal high-fat diet exposure^13,14^, while human studies have described structural and functional cardiac abnormalities from fetal life through childhood^15,16^. However, previous work has largely focused on fetal development or adult disease, leaving the postnatal maturation window underexplored. Whether maternal obesity alters the developmental trajectory of cardiomyocyte maturation itself, rather than simply triggering pathological remodeling later in life, remains unknown.

Here, we used a rat model of maternal high-fat diet exposure to investigate how maternal obesity influences postnatal cardiac maturation of offspring. By integrating longitudinal *in vivo* cardiac phenotyping with structural, transcriptomic, and metabolic analyses, we identify accelerated postnatal cardiomyocyte maturation as an early adaptive response to maternal metabolic stress. We further show that this premature maturation precedes maladaptive cardiac remodeling and ultimately predisposes the adult heart to contractile dysfunction, revealing a developmental mechanism that links maternal obesity to long-term cardiovascular vulnerability.

## METHODS

### Animal model and study design

Detailed protocols, reagents and analysis parameters are provided in the Supplemental Methods and Supplemental Table I. Wistar rats (Janvier Labs, France) were housed under specific pathogen-free conditions with ad libitum access to food and water, on a 12:12 h light/dark cycle. Female rats (mothers) were fed a normal diet (ND-Mat group) or a high-fat diet (HFD-Mat group; D12492, Research Diets, USA) for two months before mating and throughout gestation and lactation. After weaning, all offspring received a standard chow diet regardless of maternal diet. Experiments were conducted at postnatal day 5 (P5), P20, P60 and P120, depending on the analysis. Offspring of both sexes were studied, with equal numbers of males and females in each group and for every experiment. In the omics analyses, males and females were first analyzed separately. As no sex effect was found, the sexes were pooled. Dams were randomly assigned to the ND or HFD group. Animals were identified only by consecutive numbers. All experiments and analyses were performed by investigators who had access only to the animal numbers, not to the maternal diet group. No animals were excluded from the analyses. All procedures followed institutional guidelines and European Directive 2010/63/EU. The experimental protocol was approved by the French Ministry of Higher Education and Research (APAFIS #37564-2022060214594664 v17). The animal facility is authorized by the Ministère de l’Agriculture (no. 75-1090).

### In vivo cardiac and metabolic phenotyping

Echocardiography was performed under isoflurane anesthesia with a Vevo 3100 system (VisualSonics), using M-mode, Doppler, tissue Doppler imaging and speckle tracking for global longitudinal strain. Cardiac function was challenged with an intraperitoneal injection of isoproterenol (50 mg/kg). Electrocardiograms were recorded under isoflurane with needle electrodes (Emka Technologies). The P wave duration and the PR, QRS, QT and RR intervals were measured automatically, and the heart rate response to isoproterenol was followed for 4 minutes. At P60, left ventricular pressure–volume relations were recorded under ketamine/xylazine anesthesia with a 2F Millar catheter introduced through the right carotid artery. Transient inferior vena cava compression was used to derive the load-independent indices of contractility (ESPVR and PRSW) and compliance (EDPVR). Systolic blood pressure was measured by tail-cuff plethysmography (BP-2000, Bioseb) at P20 and P60.

Glucose and insulin tolerance tests (IPGTT, 2 g/kg D-glucose; IPITT, 0.75 U/kg insulin) were performed after a 6-hour fast, with tail-vein glucose measured over 120 minutes. Fat mass was measured by nuclear magnetic resonance, food intake was monitored over 2 months, and heart weight was normalized to tibia length.

### Plasma and tissue assays

Plasma glucose, triglycerides, total cholesterol and C-reactive protein were measured with an Indiko automated analyzer (Thermo Fisher Scientific). Plasma ANP and BNP were quantified at P20 and P60 by ELISA (Sigma-Aldrich), and cardiac ATP content was measured at P60 with a bioluminescence assay (Thermo Fisher Scientific).

### Cardiomyocyte isolation and mitochondrial function

Ventricular cardiomyocytes were isolated from P20 and P60 hearts by Langendorff retrograde perfusion with collagenase A. Mitochondrial respiration was measured in 20,000 permeabilized cells with an O2k high-resolution respirometer (Oroboros Instruments), using sequential additions of substrates and inhibitors to assess complex I-, II- and IV-driven respiration. Mitochondrial membrane potential was expressed as the TMRE/MitoTracker Deep Red fluorescence ratio, measured per cell by live imaging with an Operetta CLS system (PerkinElmer). For the cell death assay, cells were imaged hourly for 15 hours with TMRE and NucBlue, with or without hydrogen peroxide (10 µM) or doxorubicin (60 µM), and viability was quantified with a supervised machine-learning algorithm in Harmony software (PerkinElmer).

### Western blot and immunofluorescence

For Western blotting, proteins from frozen myocardium were separated on 10% Bis-Tris gels, transferred to nitrocellulose and probed with the antibodies listed in Supplemental Table I. Because no loading control was stable between ND-Mat and HFD-Mat hearts, protein levels were normalized to total protein (Ponceau staining). Immunofluorescence was performed on 7 µm myocardial cryosections, with DAPI and wheat germ agglutinin (WGA) to label nuclei and membranes, and imaged by deconvolution epifluorescence microscopy (DeltaVision Elite, GE Healthcare). Three-dimensional reconstructions were generated with Imaris.

### Fluorescent immunolabeling and confocal imaging of fresh cardiac biopsies

Fresh cardiac biopsies were immunolabeled and imaged using the 3D-NaissI (3D-Native Tissue Imaging) method, as previously described^17^. Briefly, fresh biopsies were incubated in HBSS with the primary antibody for 2 days at 4 °C under constant rotary agitation, then washed and incubated with the fluorescent secondary antibody, protected from light, for a further 2 days under the same conditions before imaging.

Biopsies were imaged on a Zeiss LSM 900 confocal microscope (×63 oil objective) with identical acquisition settings for a given antibody. Cardiomyocytes (CMs) were segmented with Cellpose from WGA surface staining, and a custom Fiji/ImageJ macro analyzed each CM (5 biopsies per rat, 3 stacks per biopsy, about 50–100 CMs per rat).

To quantify total expression of the protein of interest in each CM, a binary segmentation of the protein fluorescence signal was obtained by defining a threshold in the control condition for each individual experiment. Protein expression was quantified as the number of fluorescent pixels above threshold rather than as mean fluorescence intensity. This pixel-based approach was chosen because the tissue and inter-experiment variability inherent to the 3D-NaissI method precludes reliable comparison of absolute fluorescence intensities^17^.

### Histology and electron microscopy

Fibrosis was assessed on Sirius Red-stained paraffin sections and quantified with HistoLab software, and lipid accumulation on Oil Red O-stained cryosections. For transmission electron microscopy, samples were postfixed in osmium tetroxide and embedded in Epikote 812, and 90 nm sections were examined with a JEOL 1011 microscope. Mitochondrial morphology was quantified in Fiji (ImageJ). Area, circularity, aspect ratio and solidity were measured on 45 to 50 images per group and per stage, taken at ×5,000 magnification.

### Omics analyses

For RNA sequencing, total RNA was extracted from hearts at P5, P20 and P60 with TRIzol (n = 10 per group and stage; 5 males and 5 females). Stranded mRNA libraries (Illumina) were sequenced on a NovaSeq X Plus (2 × 150 bp; about 40 million reads per sample). Reads were trimmed with fastp, then aligned to the rn7 rat genome and counted with the DRAGEN Bio-IT Platform (GENCODE mRatBN7.2 annotation). Differential expression was tested with DESeq2. Genes with an absolute log₂ fold change above 1 and a Benjamini–Hochberg false discovery rate below 0.05 were considered differentially expressed. Gene set enrichment analysis used clusterProfiler and the KEGG database.

Untargeted metabolomics of heart tissue and plasma was performed by liquid chromatography–high-resolution mass spectrometry (Q Exactive, Thermo Fisher Scientific; ZIC-pHILIC column). Free fatty acids were profiled by gas chromatography–mass spectrometry after pentafluorobenzyl bromide derivatization (both: n = 6 per group and stage; 3 males and 3 females). Metabolites were annotated with HMDB and KEGG. Data deposition is described in the Data Availability section.

### Statistics

Statistical analyses were performed with GraphPad Prism (version 11, GraphPad Software). Data are presented as mean ± SD unless otherwise stated. Normality of the data was checked with the Shapiro-Wilk test before each statistical test was applied. Two groups were compared with an unpaired two-tailed Student’s t-test. For more than two groups, one-way or two-way ANOVA was followed by Tukey’s post-hoc test; three-way ANOVA (Supplemental Figures 6 and 8) and Pearson’s correlation were used where indicated in the legends. Sex was tested as a factor in the omics analyses. No sex effect was found, so male and female data were pooled. Differences were considered significant at P<0.05 (*P<0.05, **P<0.01, ***P<0.001, ****P<0.0001).

## RESULTS

### Maternal high-fat diet induces postnatal growth abnormalities and biphasic cardiac functional remodeling in the offspring (Figure 1)

Whether maternal obesity alters the trajectory of postnatal cardiac growth and function, rather than simply predisposing to disease later in life, remains unclear. To address this, offspring born to females fed a normal (ND-Mat) or high-fat (HFD-Mat) diet were longitudinally phenotyped across four postnatal stages spanning infancy to adulthood (P5, P20, P60 and P120) (Figure 1A). HFD-Mat offspring exhibited significantly increased body weight and heart weight compared with ND-Mat offspring at all developmental stages examined (Figure 1B, 1C). Food intake was monitored to determine whether increased body weight in HFD-Mat offspring was associated with hyperphagia. No difference in food consumption was observed between ND-Mat and HFD-Mat offspring, indicating that the increased body weight of HFD-Mat offspring cannot be attributed to increased caloric intake (Supplemental Figure 1A). Body composition analysis by nuclear magnetic resonance (NMR) further revealed a significant increase in fat mass in HFD-Mat offspring (Figure 1D).

**Figure 1:**
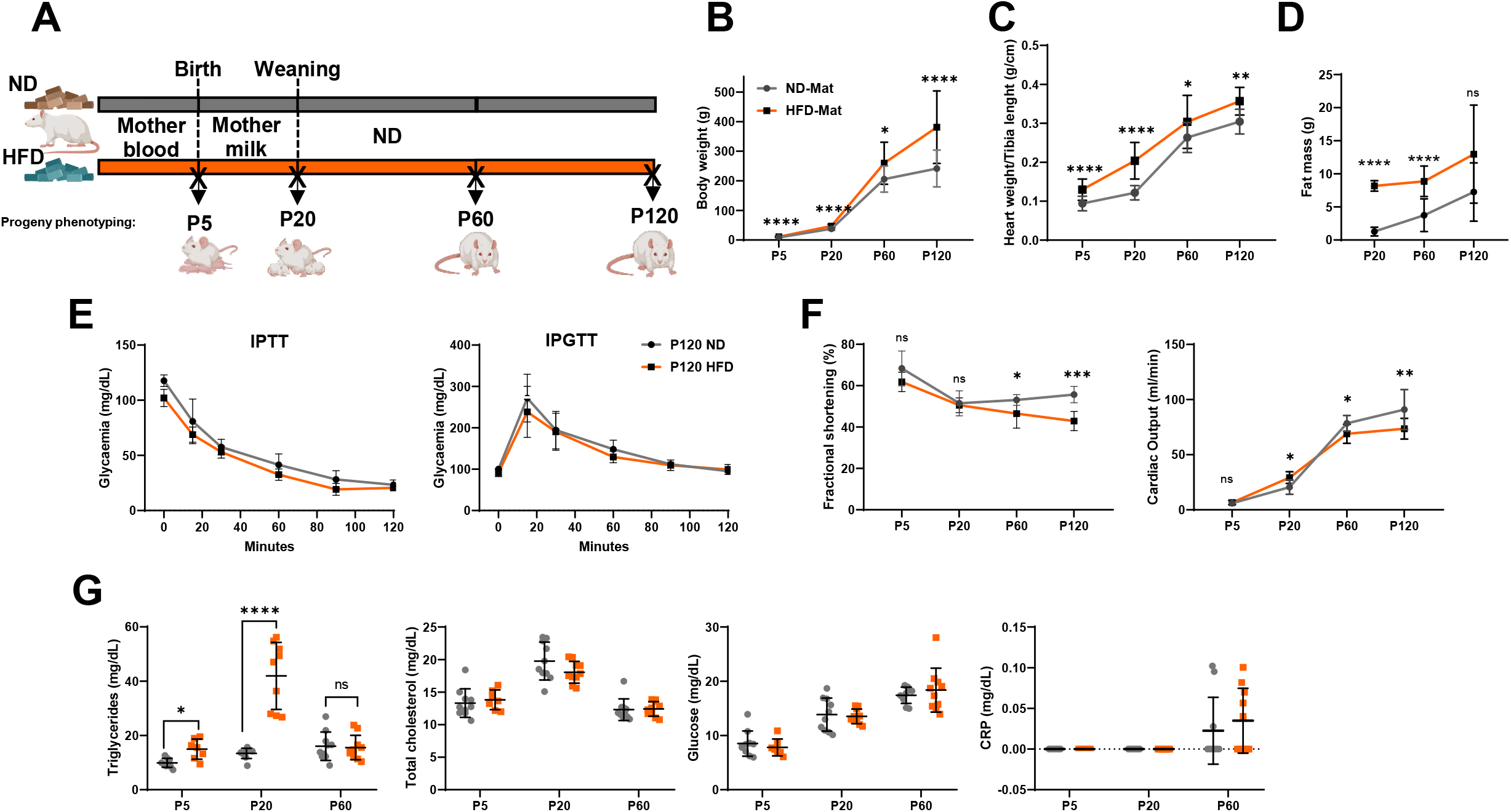
Maternal high-fat diet induces postnatal growth abnormalities and biphasic cardiac functional remodeling in the offspring. **(A)** Schematic of experimental design. **(B)** Body weight measurements of offspring from normal diet (ND-Mat, grey) and high-fat diet (HFD-Mat, orange) mothers at postnatal days 5 (P5), 20 (P20), 60 (P60) and 120 (P120) (N=8 to 30 rats/group). **(C)** Heart weight was normalized to tibia length (N=8 to 34 rats/group). **(D)** Fat mass was assessed by nuclear magnetic resonance (NMR) at P20, P60 and P120 in offspring from ND and HFD mothers (N=6 to 8 rats/group). **(E)** Intraperitoneal glucose and insulin tolerance tests (IPGTT and IPITT, respectively) were performed in both groups at P120 (N=6 to 8 rats/group). **(F)** Systolic function parameters assessed by echocardiography in offspring including fractional shortening (FS) and cardiac output (CO) (N=8 rats/group). **(G)** Blood levels of triglycerides, total cholesterol, glucose and CRP (in mg/dL) were quantified using the INDIKO analyzer. Data are presented as mean±SD; two-way ANOVA followed by Tukey’s multiple comparisons test; *P<0.05; **P<0.01; ***P<0.001; **** P<0.0001; ns, not significant.

To determine whether maternal HFD induced systemic metabolic alterations in the offspring, intraperitoneal glucose tolerance tests (IPGTT) and insulin tolerance tests (IPITT) were performed. HFD-Mat offspring exhibited normal glucose tolerance and insulin sensitivity, ruling out overt glucose intolerance or insulin resistance (Figure 1E). Systolic blood pressure, measured by tail-cuff plethysmography, was unchanged between ND-Mat and HFD-Mat offspring at both P20 and P60, indicating that maternal HFD-Mat does not induce hypertension during postnatal development (Supplemental Figure 1B).

Cardiac function was next assessed by echocardiography. Fractional shortening was significantly reduced at the later postnatal stages (P60 and P120) in HFD-Mat offspring (Figure 1F, Supplemental Table 1A, 1B). In contrast, cardiac output was significantly increased at P20 before declining at P60 and P120 (Figure 1F), suggesting an early compensatory response followed by progressive functional deterioration in HFD-Mat offspring.

Circulating triglyceride, total cholesterol, glucose and C-reactive protein (CRP) concentrations were next quantified in plasma using a clinical chemistry analyzer. Triglyceride levels were elevated at P5 and P20 in HFD-Mat offspring before returning to control levels by P60, whereas total cholesterol and CRP concentrations remained unchanged across all time points (Figure 1G).

Collectively, these results show that maternal HFD-Mat reprograms offspring growth and body composition and drives a biphasic trajectory of cardiac function — an early, unexpected gain in cardiac output followed by a later decline in systolic performance.

### Maternal high-fat diet induces early cardiac adaptation followed by progressive functional decline and concentric remodeling (Figure 2)

This biphasic trajectory raised the question of whether the early functional gain in HFD-Mat offspring reflects genuine adaptation or an early sign of pathological remodeling. Comprehensive cardiac phenotyping was therefore performed at P20 and P60 (Supplemental Table 1A).

**Figure 2:**
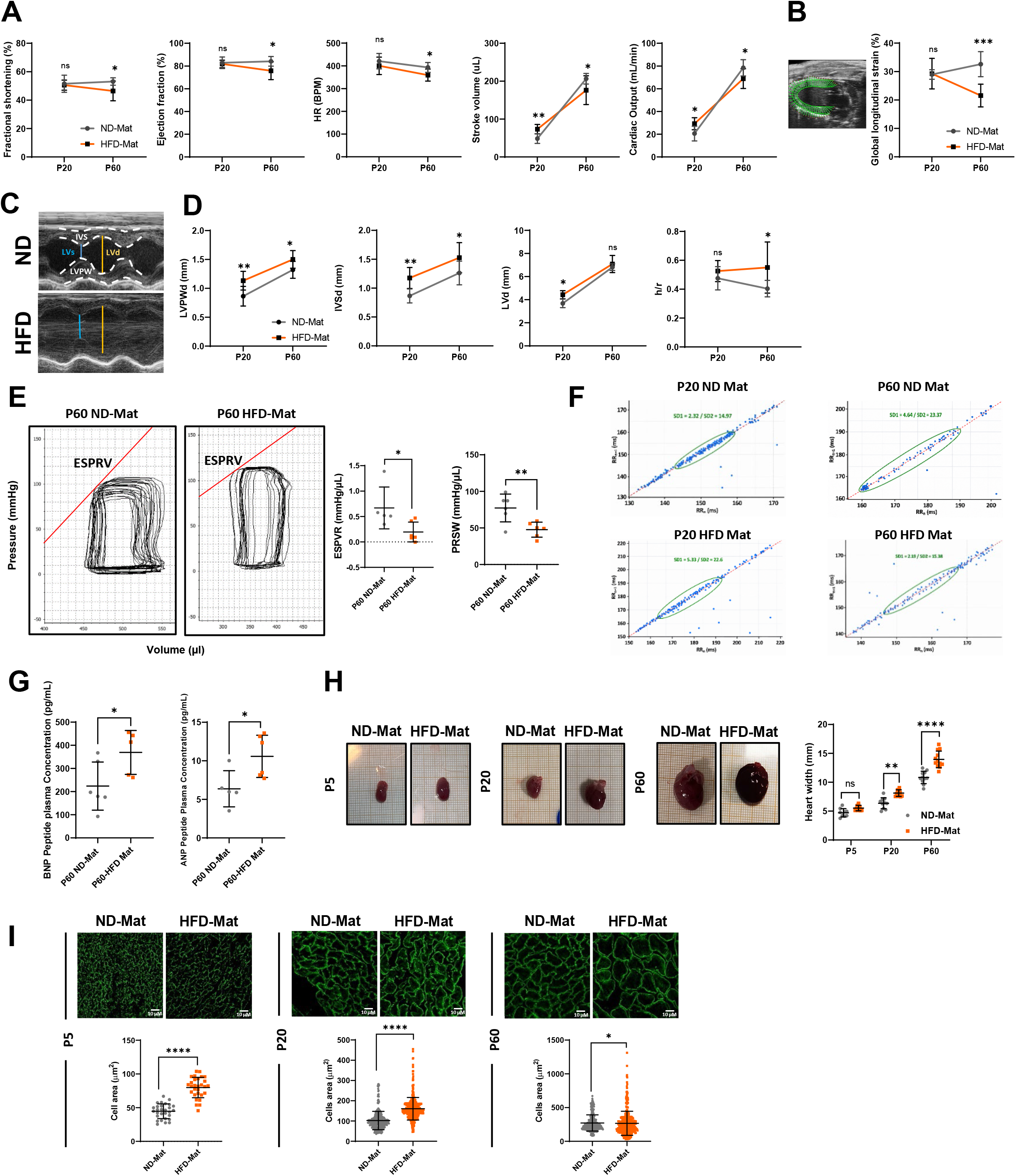
Maternal high-fat diet induces early cardiac adaptation followed by progressive functional decline and concentric remodeling. **(A)** Systolic function assessed by echocardiography on P20 and P60 offspring born to ND (ND-Mat, grey) and HFD (HFD-Mat, orange) mothers, including fractional shortening (FS), ejection fraction (EF), heart rate (HR), stroke volume (SV), and cardiac output (CO) (N=8 rats/group). **(B)** Myocardial contractility measured by global longitudinal strain (GLS) (N=8 rats/group). **(C)** Representative M-mode echocardiographic images showing left ventricular internal diameters in diastole (LVd) and systole (LVs), interventricular septum (IVS), and left ventricular posterior wall (LVPW). **(D)** Morphometric parameters: left ventricular posterior wall thickness in diastole (LVPWd), interventricular septum thickness in diastole (IVSd), and relative wall thickness (h/r, calculated as the ratio of (LVPWd + IVSd) to LV internal diameter) (N=8 rats/group). **(E)** Representative pressure-volume loops of P60 ND-Mat and HFD-Mat offspring and quantification of systolic function parameters (end systolic pressure volume relationship (ESPVR) and preload recruitable stroke work (PRSW)) (N=6 rats/group)**. (F)** Poincaré plot representation of consecutive RR intervals, defined as the time intervals between two successive R-wave peaks on the electrocardiogram (ECG), at P20 and P60 in both groups (N = 8 rats/group). **(G)** Plasma concentrations of B-type natriuretic peptide (BNP) and atrial natriuretic peptide (ANP) measured by ELISA (N=5 to 6 rats/group). **(H)** Heart width measured at postnatal days P5, P20, and P60 (N=9 to 12 rats/group). **(I)** Representative immunofluorescence images of transverse heart tissue sections at the three developmental stages, with dystrophin (green) as a membrane marker (scale bar 10 µm); cardiomyocyte cross-sectional area was quantified (N=6 rats/group). Data are presented as mean±SD; two-way ANOVA followed by Tukey’s multiple comparisons test (A, B, D, F, H) and Unpaired Student’s *t*-test (E, G, I); \**P<*0.05, \*\**P*<0.01; \*\*\**P*<0.001, \*\*\*\**P*<0.0001; ns, not significant.

At P20, HFD-Mat offspring exhibited increased stroke volume and cardiac output without changes in systolic function, consistent with an early compensatory phase (Figure 2A). By contrast, adult (P60) HFD-Mat offspring showed significantly reduced ejection fraction, fractional shortening, heart rate, stroke volume and cardiac output (Figure 2A), alongside reduced myocardial contractility, reflected by decreased global longitudinal strain (GLS) at P60 (Figure 2B) and further reduced at P120 (Supplemental Figure 1D). HFD-Mat offspring also showed an enhanced stroke volume and cardiac output response to isoproterenol at P20, an effect no longer present at P60 (Supplemental Figure 2). Morphometric analysis further revealed increased LVPW and IVS thickness without changes in LV diameter, consistent with concentric hypertrophy already detectable at P20 (Figure 2C, 2D, Supplemental Figure 1C).

Diastolic function was largely preserved early on: E/A and E/E′ ratios were reduced at P20 but normalized by P60, with only modest IVRT prolongation (Supplemental Figure 3A). By P120, however, E/E′ — which continued to physiologically decline in ND-Mat offspring — remained stable in HFD-Mat offspring, resulting in a significant elevation preceded by an early, sustained rise in left atrial area already present at P20 (Supplemental Figure 3B).

Invasive pressure-volume loop analysis at P60 confirmed impaired load-independent contractility, with reduced ESPVR and PRSW (Figure 2E), while EDPVR was unchanged, indicating preserved passive compliance (Supplemental Figure 3C).

Poincaré plot analysis of ECG-derived RR intervals showed a broader ellipse at P20 in HFD-Mat offspring (increased SD1/SD2, greater beat-to-beat variability), which reversed by P60 into a markedly narrower ellipse (reduced SD1/SD2 despite the physiological increase seen in ND-Mat offspring) — consistent with altered autonomic modulation in adulthood (Figure 2F).

Circulating ANP and BNP were significantly increased in P60 HFD-Mat offspring (Figure 2G), alongside increased heart width from the earliest postnatal stages (Figure 2H) and increased cardiomyocyte cross-sectional area from early life onward (Figure 2I).

Despite this hypertrophy and systolic dysfunction, histological analyses at P60 showed no evidence of fibrosis (Sirius Red, Supplemental Figure 4A) or lipid accumulation (Oil Red O, Supplemental Figure 4B; TEM lipid droplet content, Supplemental Figure 4C), indicating that the phenotype is not driven by direct lipotoxic injury and is more consistent with developmentally programmed metabolic remodeling.

Overall, maternal HFD-Mat triggers an early compensatory cardiac phase that progresses to systolic dysfunction, impaired contractility and concentric hypertrophy by adulthood.

### Early-life metabolic stress reshapes the developmental trajectory of the cardiac transcriptome (Figure 3)

To identify the molecular programs underlying this early adaptation and its later failure, bulk RNA sequencing was performed on whole hearts from ND-Mat and HFD-Mat offspring across the P5-P20-P60 window (Figure 3A).

**Figure 3:**
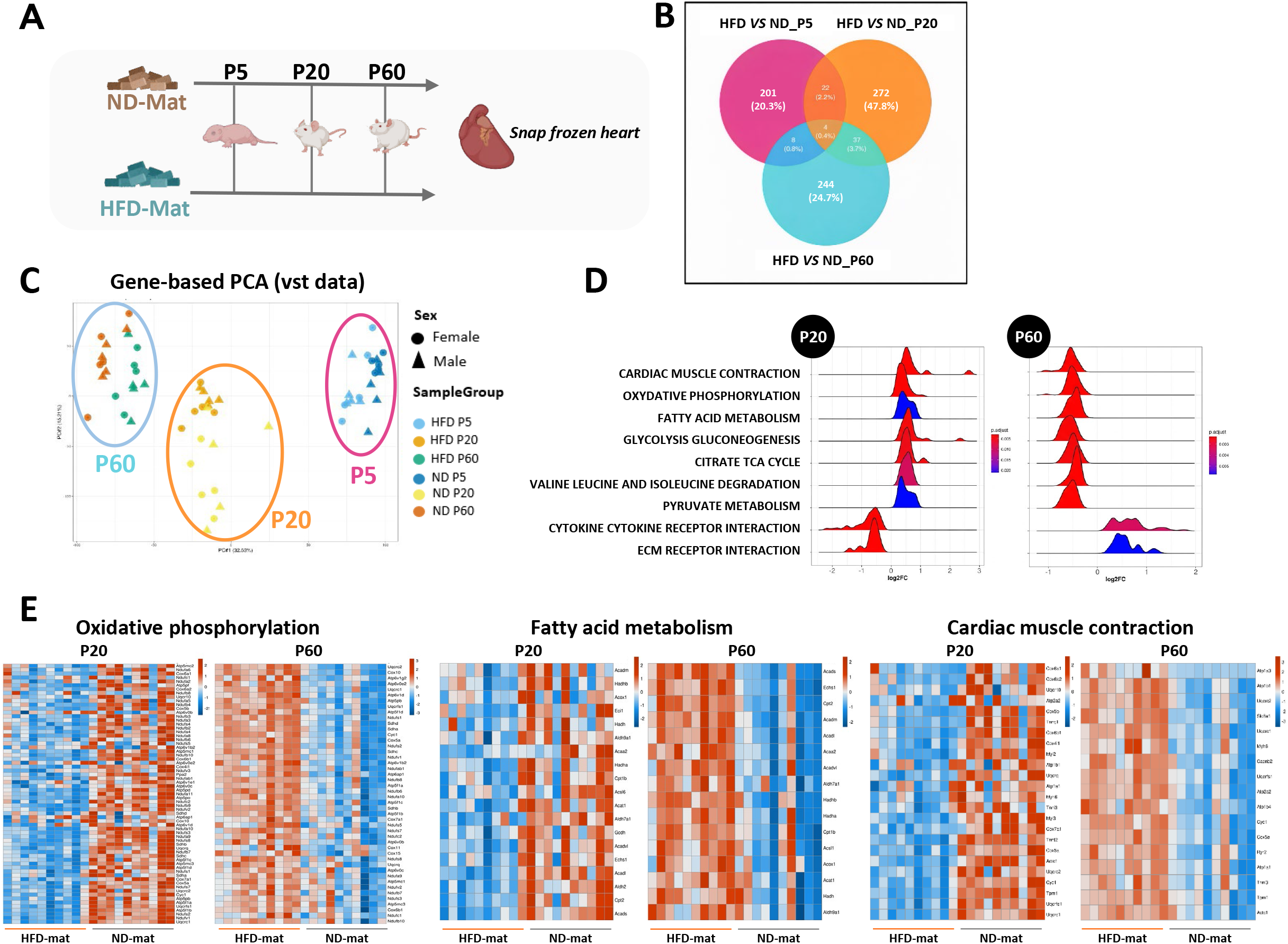
Early-life metabolic stress reshapes the developmental trajectory of the cardiac transcriptome. **(A)** Schematic of the experimental protocol (N=10 rats/group). **(B)** Venn diagram of differentially expressed genes between ND-Mat and HFD-Mat offspring at three developmental stages in offspring (pink: P5, orange: P20, blue: P60). **(C)** Principal component analysis (PCA) showing the distribution of samples according to sex, based on variance-stabilizing transformed (VST) RNA-seq count data. **(D)** Ridge plot of the top regulated pathways in HFD-Mat versus ND-Mat offspring. **(E)** Heatmap of three pathways: oxidative phosphorylation, fatty acid metabolism, and cardiac contraction at P20 and P60, with blue indicating downregulation and red upregulation.

Differential expression analysis identified marked transcriptomic remodeling at each developmental stage, with 201, 472, and 244 differentially expressed genes (DEGs) detected at P5, P20, and P60, respectively, as illustrated by the Venn diagram (Figure 3B). Principal component analysis (PCA) revealed a clear segregation according to developmental stage, whereas no sex-dependent differences were observed within each group (Figure 3C).

KEGG pathway enrichment analysis revealed a striking stage-dependent pattern of pathway regulation as shown by the ridge plot (Figure 3D).

At P20, genes involved in cardiac muscle contraction, fatty acid metabolism, and oxidative phosphorylation were significantly enriched among upregulated pathways in HFD-Mat offspring. In contrast, these same pathways were significantly downregulated at P60 (Figure 3D). Heatmap visualization confirmed this biphasic regulation, highlighting coordinated changes in genes associated with cardiac contraction, oxidative phosphorylation, and fatty acid metabolism (Figure 3E). This biphasic transcriptional signature closely paralleled the functional trajectory observed in vivo, with early upregulation of contractile and metabolic genes coinciding with the compensatory increase in stroke volume and cardiac output at P20, and their subsequent downregulation mirroring the decline in contractility and systolic function observed at P60. Notably, the downregulation of fatty acid oxidation and oxidative phosphorylation genes alongside impaired contractile function is reminiscent of the metabolic remodeling classically described in pathological cardiac hypertrophy and heart failure^18^, suggesting that HFD-Mat offspring recapitulate a conserved molecular signature of cardiac stress independent of maternal diet exposure.

Together, these findings indicate that maternal HFD-Mat does not simply perturb the cardiac transcriptome, but rather induces a temporal shift in its regulation, consistent with premature activation and subsequent premature repression of contractile and metabolic gene programs.

### Early-life metabolic stress selectively remodels cardiac lipid metabolism during postnatal development (Figure 4)

To test whether the transcriptional induction of fatty-acid metabolism genes translated into altered lipid handling, lipidomic and metabolomic profiling of heart and plasma was performed across the same developmental window (Figure 4A; no sex-dependent clustering was observed, Supplemental Figure 5A).

**Figure 4:**
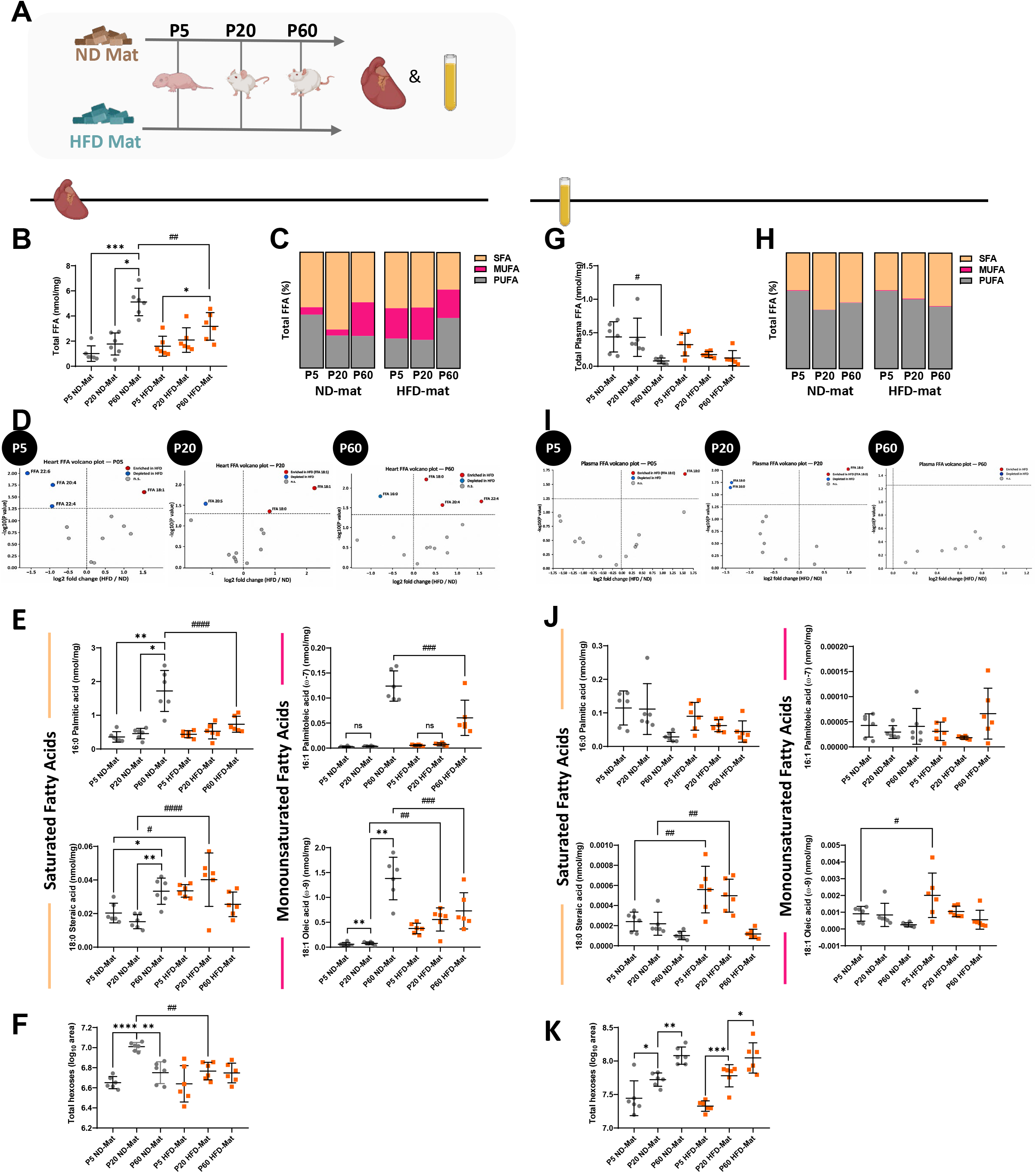
Early-life metabolic stress selectively remodels cardiac lipid metabolism during postnatal development. **(A)** Schematic of the experimental protocol (N=6 rats/group). **(B)** Total free fatty acid (FFA) levels (expressed in nmol/mg) at the three developmental stages in both groups, in heart and **(G)** plasma. **(C)** Distribution of the three FFA classes, saturated fatty acids (SFA), monounsaturated fatty acids (MUFA) and polyunsaturated fatty acids (PUFA), in heart and **(H)** plasma, expressed as percentages. **(D)** Volcano plots showing differential metabolite abundance between HFD-Mat and ND-Mat offspring in the **heart** and **plasma (I)** at P5, P20 and P60. The x-axis represents the log2 fold change (HFD-Mat/ND-Mat), and the y-axis represents the −log10 P value. Metabolites significantly enriched in HFD-Mat offspring are shown in red, whereas significantly depleted metabolites are shown in blue; non-significant metabolites are shown in grey. **(E)** Representative levels of SFA (palmitic acid and stearic acid) and MUFA (palmitoleic acid and oleic acid) in each experimental group in heart and **(J)** plasma (expressed in nmol/mg). **(F)** Graph representing total hexoses (expressed as −log10 area) at the three stages in both groups, in heart and **(K)** plasma. Data are presented as mean±SD; two-way ANOVA followed by Tukey’s multiple comparisons test; \**P*<0.05, \*\**P*<0.01, \*\*\**P*<0.001, \*\*\*\**P*<0.0001, difference within a group (developmental effect); ^#^*P*<0.05, ^##^*P*<0.01, ^###^*P*<0.001, ^####^*P*<0.0001, difference between groups (treatment effect).

Cardiac lipidomic profiling revealed a marked reduction in total free fatty acid (FFA) content in HFD-Mat offspring at P60 (Figure 4B). While total cardiac FFAs increased markedly between P20 and P60 in ND controls, reflecting normal postnatal maturation of lipid metabolism, this developmental increase was absent in HFD-Mat offspring, indicating an altered trajectory of cardiac FFA accumulation (Figure 4B). Analysis of FFA subclasses revealed a stage-dependent remodeling, with monounsaturated fatty acids (MUFAs) enriched at P5 and P20 but reduced at P60, and saturated fatty acids (SFAs) consistently reduced throughout postnatal development (Figure 4C). This was driven at the individual lipid level by reductions in palmitic and stearic acid (SFAs) and in palmitoleic and oleic acid (MUFAs) at P60, despite a transient increase in stearic acid at P20 (Figure 4D, 4E).

These alterations were largely restricted to the heart: plasma lipidomic analysis revealed no differences in total circulating FFAs or FFA subclass distribution between groups (Figure 4G, 4H), although stearic acid was transiently increased in plasma at P20 and P60, and oleic acid at P5 (Figure 4I, 4J). Metabolomic analysis identified only limited additional alterations, with cardiac hexoses specifically reduced at P20 (Figure 4F) while circulating hexose levels remained unchanged (Figure 4K). This reduction coincided with a significant upregulation of the glycolysis/gluconeogenesis pathway in the P20 cardiac transcriptome (Figure 3D, Supplemental Figure 5B), in line with increased cardiac glucose utilization during the early compensatory phase. Consistent with this tissue-specific pattern, diet × developmental stage interaction effects on individual fatty acid species were markedly stronger in the heart than in plasma (Supplemental Figure 5C).

Collectively, these findings indicate that maternal HFD-Mat reprograms the developmental trajectory of cardiac, but not circulating, fatty acid accumulation, in a tissue-specific manner not mirrored by the circulating lipid metabolome.

### Cardiac mitochondrial homeostasis is altered during postnatal development in HFD-Mat offspring (Figure 5)

Given that mitochondria govern cardiac lipid and energy metabolism, we next assessed mitochondrial structure and function at P60, the stage at which cardiac dysfunction was most pronounced.

**Figure 5:**
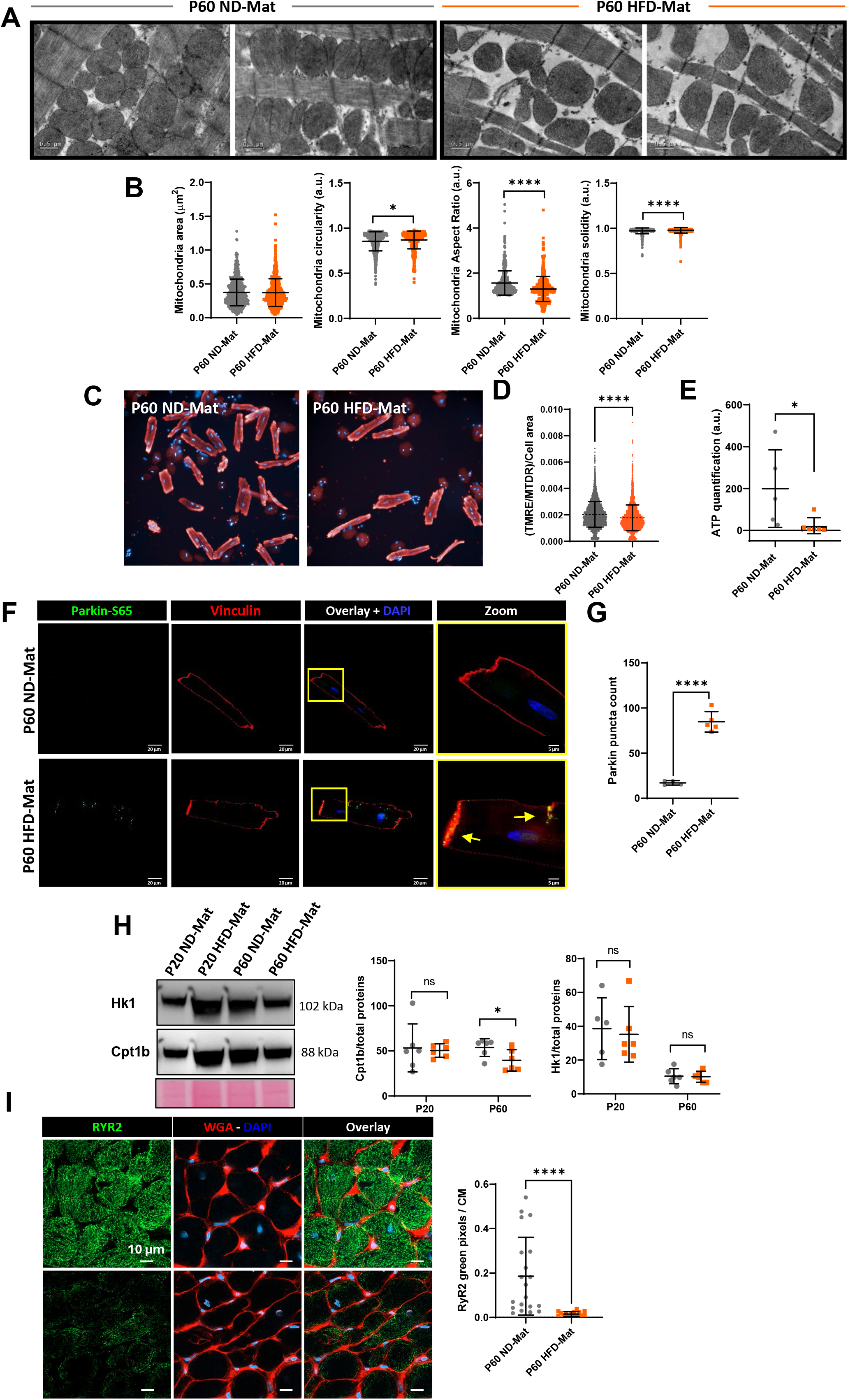
Cardiac mitochondrial homeostasis is altered during postnatal development in HFD-Mat offspring. **(A)** Representative transmission electron microscopy images (1,500× magnification). White arrows indicate mitochondria (N=4 rats/group). **(B)** Quantification of mitochondrial area, circularity, aspect ratio (AR) and solidity at the P60 stage. **(C)** Representative images of isolated adult cardiomyocytes (P60) stained with tetramethylrhodamine ethyl ester (TMRE) and MitoTracker Deep Red (MTDR) to assess mitochondrial membrane potential. **(D)** Quantification of the TMRE/MTDR fluorescence ratio normalized to cell area (N=4 rats/group). **(E)** Quantification of adenosine triphosphate (ATP) content in cardiac tissue at P60 using a luminescence-based ATP assay kit (N=6 rats/group). **(F)** Representative immunofluorescence images of isolated P60 cardiomyocytes stained for phosphorylated Parkin at serine 65 (Parkin S65) in green (scale bar, 20 µm; zoom, 5 µm). **(G)** Quantification of Parkin S65-positive puncta (N=4 to 5 rats/group). **(H)** Representative immunoblots and corresponding quantification of the relative protein expression of hexokinase 1 (HK1) and carnitine palmitoyltransferase 1B (CPT1B), normalized to total protein staining (Ponceau S) (N=6 rats/group). **(I)** Representative immunofluorescence images showing CPT1B staining (in green) and quantification in fresh-frozen cardiac biopsies from ND-Mat and HFD-Mat offspring at P60 (N=3 rats/group). Data are presented as mean±SD; two-way ANOVA followed by Tukey’s multiple comparisons test (H) and Unpaired Student’s *t*-test (B, D, E, G, I); \**P*<0.05; \*\**P*<0.01; \*\*\**P*<0.001; **** *P*<0.0001; ns, not significant.

Transmission electron microscopy revealed that mitochondrial size did not differ between HFD-Mat offspring and ND-Mat controls; however, HFD-Mat mitochondria displayed significantly increased circularity and solidity, together with a reduced aspect ratio, indicating a rounder and more compact mitochondrial shape (Figure 5A, 5B). To determine whether these morphological changes were associated with altered function, we measured mitochondrial membrane potential in primary adult cardiomyocytes using the potentiometric dye tetramethylrhodamine ethyl ester (TMRE), normalizing to mitochondrial content with MitoTracker DeepRed (MTDR) and to cell size. We observed a reduced TMRE/MTDR/Cell area ratio, indicating a decrease in mitochondrial membrane potential (Figure 5C, 5D), which was accompanied by reduced cardiac ATP content in the HFD-Mat group (Figure 5E). Puncta quantification of phospho-Parkin (Ser65), a marker of dysfunctional mitochondria, revealed a significant increase in Parkin Ser65-positive puncta in HFD-Mat cardiomyocytes (Figure 5F, 5G).

RNA sequencing further revealed a biphasic transcriptional regulation of oxidative phosphorylation complex subunits in HFD-Mat hearts: transcripts encoding Ndufs1 and *Ndufv1* (complex I), *Sdhb* (complex II), Uqcrc2 (complex III), and *Cox5* (complex IV), together with *Ldha* (glycolysis) and *Acadl* (fatty acid β-oxidation), were significantly increased at P20 but subsequently decreased at P60 relative to ND controls (Supplemental Figure 6A).

Despite these structural and transcriptional alterations, HFD-Mat hearts showed no differences in OXPHOS protein expression, mitochondrial respiratory capacity, or cardiomyocyte survival following H₂O₂ exposure, indicating that maternal HFD-Mat does not compromise global mitochondrial integrity or increase susceptibility to cell death (Supplemental Figure 6B-D).

Western blot analysis revealed a significant reduction in the expression of carnitine palmitoyl transferase 1B (CPT1B), the rate-limiting enzyme controlling mitochondrial long-chain fatty acid import and β-oxidation, in P60 HFD-Mat hearts. However, despite reaching statistical significance, this reduction was relatively modest (∼15%), suggesting a moderate rather than a major alteration in CPT1B expression. In contrast, the expression of hexokinase 1 (HK1), a key enzyme initiating the glycolytic pathway through glucose phosphorylation, remained unchanged (Figure 5H). Consistent with these findings, immunofluorescence performed on fresh cardiac biopsies demonstrated a marked decrease in CPT1B labelling in the HFD-Mat group at P60 (Figure 5I).

Collectively, these data indicate that maternal HFD-Mat spares global mitochondrial respiratory capacity and OXPHOS protein content in the offspring heart, while inducing mitochondrial morphological remodeling, reduced membrane potential, and a moderate reduction in CPT1B expression at P60.

### Maternal HFD-Mat accelerates early sarcomere maturation but impairs the adult contractile apparatus (Figure 6)

The transcriptomic enrichment of the cardiac muscle contraction pathway at P20 prompted us to investigate whether maternal HFD-Mat alters the structural maturation of the sarcomere (Figure 6A). RNA sequencing revealed an increased expression of key sarcomeric structural genes, including *Acta1*, encoding α-skeletal actin, and *Actn2*, encoding α-actinin-2, a major Z-disc protein responsible for anchoring actin filaments and maintaining sarcomere integrity, *Tpm1*, encoding tropomyosin 1, a thin-filament protein regulating actin-myosin interaction, and *Tnnt2*, encoding cardiac troponin T, a core component of the troponin complex controlling calcium-dependent contraction. All four transcripts were significantly upregulated at the juvenile stage (P20), consistent with an accelerated activation of the contractile program, but were subsequently downregulated in adulthood (P60), together with a reduced Myh6/Myh7 ratio, suggesting a decline in sarcomeric gene expression by adulthood (Figure 6B). Consistent with this transcriptional decline, immunoblot analysis confirmed a significant decrease in cardiac troponin T (TNNT2) protein expression at P60 in HFD-Mat offspring (Figure 6C).

**Figure 6:**
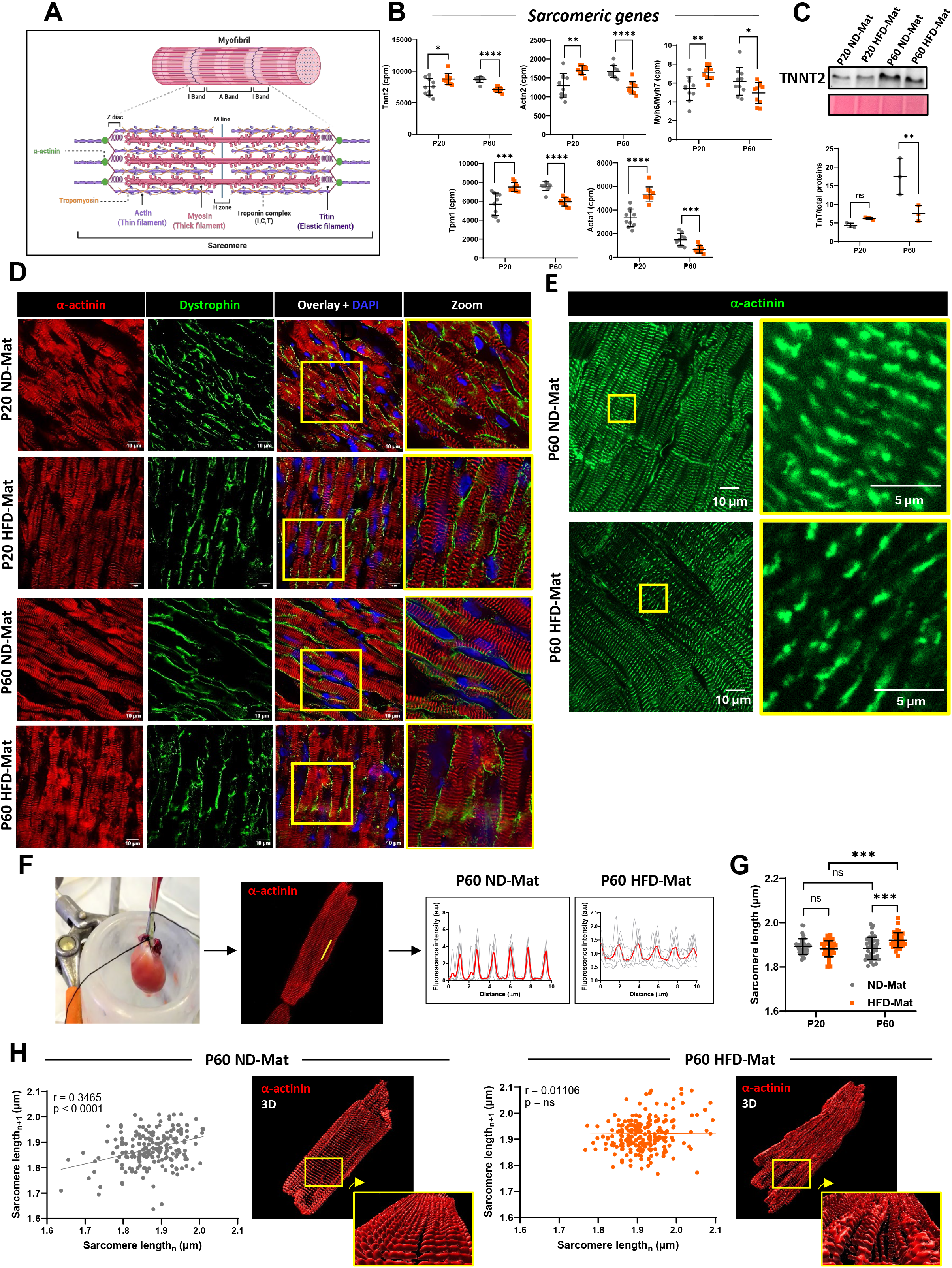
Maternal HFD accelerates early sarcomere maturation but promotes maladaptive sarcomeric remodeling in adulthood. **(A)** Schematic representation of the sarcomere and its major structural components. **(B)** Normalized RNA sequencing expression (counts per million, CPM) of the sarcomeric genes *Tnnt2*, *Actn2*, *Myh6/Myh7* ratio, *Tpm1* and *Acta1* at P20 and P60 stages (N=10 rats/group). **(C)** Representative immunoblot and quantification of TNNT2 (TnT) protein expression at P20 and P60 stages (N=3 rats/group). **(D)** Representative immunofluorescence images of longitudinal heart sections stained for α-actinin-2 (ACTN2, in red) to visualize sarcomere organization at P20 and P60 stages in ND-Mat and HFD-Mat offspring (N=6 rats/group). **(E)** Representative immunofluorescence images showing α-actinin (in green) staining and quantification in fresh-frozen cardiac biopsy sections from ND-Mat and HFD-Mat offspring at P60 (N=3 rats/group). **(F)** Schematic of Langendorff-based cardiomyocyte isolation from P20 and P60 hearts, and frequency distribution of α-actinin-2 striation periodicity in isolated cardiomyocytes from ND-Mat and HFD-Mat offspring. **(G)** Quantification of sarcomere length in isolated cardiomyocytes, measured by tracing a line perpendicular to consecutive sarcomeric striations (N=3 rats/group). **(H)** Correlation analysis of sarcomere length measurements illustrating sarcomeric organization in ND-Mat and HFD-Mat offspring at P60 (N=3 rats/group, Pearson’s correlation analysis) and representative 3D volume-rendered images of sarcomere structure. Data are presented as mean±SD; two-way ANOVA followed by Tukey’s multiple comparisons test (B, C, G) and Pearson’s correlation analysis (H); \**P*<0.05, \*\**P*<0.01, \*\*\**P*<0.001, **** *P*<0.0001; ns, not significant.

To determine whether these molecular changes translated into structural remodeling, sarcomere organization was examined by α-actinin immunofluorescence on longitudinal cardiac tissue sections. At P20, HFD-Mat hearts displayed a more regular and organized striated pattern than ND-Mat controls, suggesting accelerated maturation of the contractile apparatus during early postnatal life (Figure 6D). In contrast, this organized architecture was no longer maintained at P60, with HFD-Mat cardiomyocytes exhibiting a marked disorganization of the striated pattern. Consistent with this, immunofluorescence performed on fresh cardiac biopsies at P60 confirmed a pronounced disruption of α-actinin organization in HFD-Mat hearts, characterized by Z-discs misalignment and a near-complete loss of the regular striation compared with controls (Figure 6E).

To further assess sarcomere structural integrity at the cellular level, cardiomyocytes were isolated from Langendorff-perfused hearts and immunostained for α-actinin to visualize the Z-disc lattice. Sarcomere length did not differ between ND-Mat and HFD-Mat cardiomyocytes at P20, but became significantly increased in HFD-Mat cardiomyocytes by P60, although values remained within the physiological range (1.7-1.9 µm at rest^19^) (Figure 6F, 6G). Isolated HFD-Mat cardiomyocytes were already significantly longer and wider than ND-Mat controls at P20, and this difference persisted at P60, indicating that maternal HFD-Mat induces early and sustained cardiomyocyte hypertrophy preceding sarcomere lengthening (Supplemental Figure 7A). Sarcomere length did not correlate with cardiomyocyte size in either group at either age, indicating that these sarcomere-level changes were independent of overall cell size (Supplemental Figure 7B).

To determine whether the age-dependent lengthening observed in HFD-Mat cardiomyocytes was accompanied by a loss of structural organization, we next examined the relationship between the lengths of consecutive sarcomeres along individual myofibrils at P60. In ND-Mat cardiomyocytes, the length of a given sarcomere (position *n*) was significantly correlated with that of the immediately adjacent sarcomere (*n+1*). In other words, neighboring sarcomeres tended to have similar lengths, indicating a regular and coordinated organization of sarcomeres along the myofibril. Strikingly, this correlation was completely abolished in HFD-Mat cardiomyocytes (Figure 6H). Consistently, 3D reconstruction of α-actinin labelling confirmed a well-aligned, parallel Z-disc array in ND-Mat cardiomyocytes. This contrasted with a visibly disorganized myofibrillar lattice in HFD-Mat cells (Figure 6H).

Together, these findings indicate that maternal HFD-Mat drives early cardiomyocyte hypertrophy followed by progressive sarcomeric remodeling, combining structural elongation with a loss of spatial coordination between neighboring sarcomeres at P60.

### Progressive remodeling of sarcomeric and excitation-contraction coupling proteins in adult HFD-Mat offspring (Figure 7)

Given the structural remodeling of the sarcomere observed in P20 and P60 HFD-Mat hearts, we next investigated whether proteins involved in excitation-contraction coupling and membrane organisation (Figure 7A) were similarly affected. At P20, transcripts encoding key calcium-handling proteins, including *Jph2* (Junctophilin-2), *Ryr2*, *Atp2a2* (SERCA2a) and *Cav3*, were significantly upregulated, consistent with the early compensatory activation of the contractile program described above. At P60, however, transcriptomic analysis revealed a coordinated downregulation of these same calcium-handling genes, together with *Cav3*, encoding caveolin-3, the muscle-specific structural component of caveolae required for sarcolemmal and T-tubule organization (Figure 7B).

**Figure 7.**
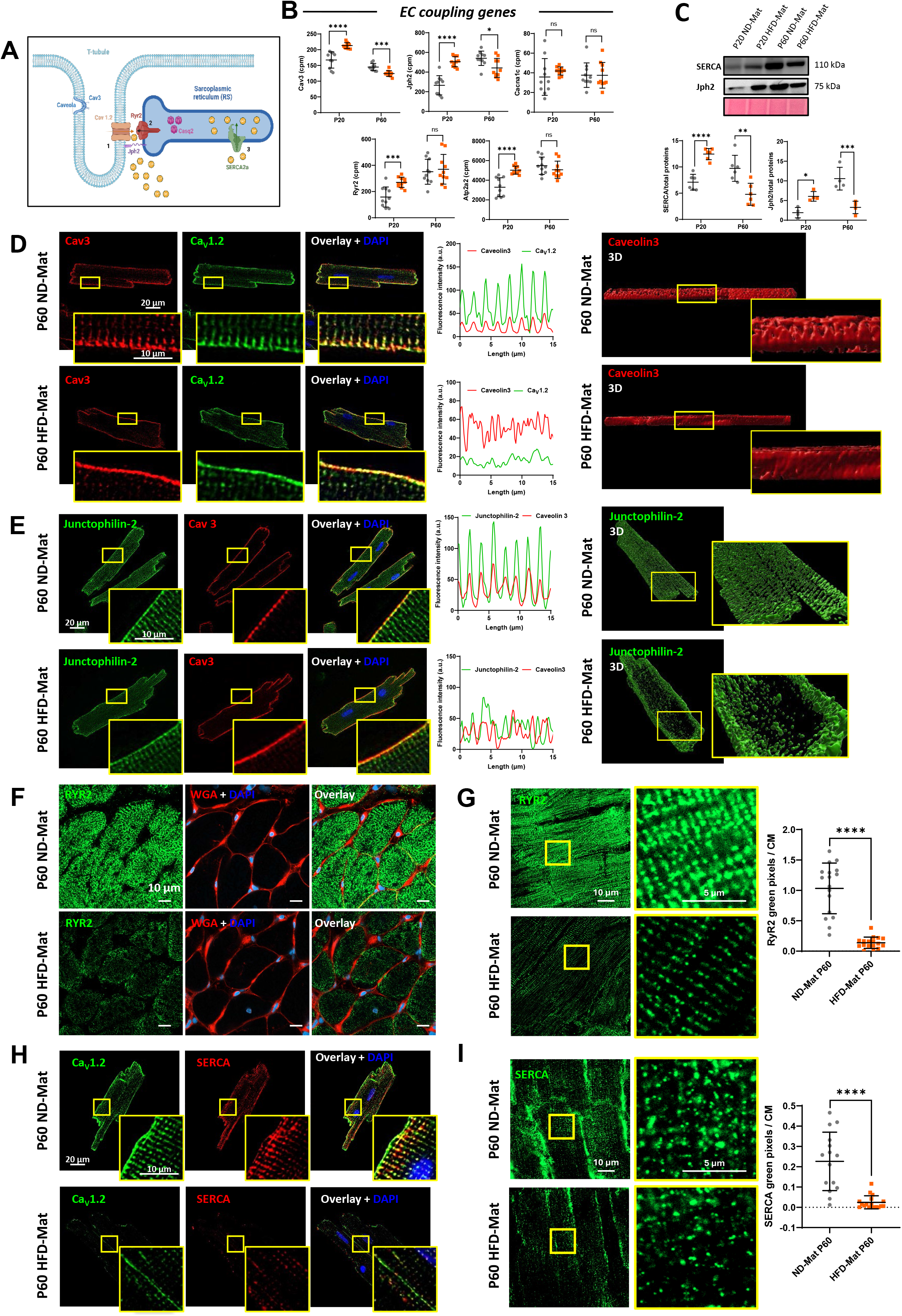
Progressive remodeling of the contractile machinery extends to excitation-contraction coupling proteins. **(A)** Schematic representation of the cardiac T-tubule/sarcoplasmic reticulum couplon, illustrating the spatial organization of caveolin-3 (Cav3), junctophilin-2 (JPH2), the L-type calcium channel (Cav1.2), the ryanodine receptor 2 (RyR2), and SERCA2a. **(B)** Normalized RNA sequencing expression (counts per million, CPM) of *Cav3*, *Jph2*, *Cacna1c*, *Ryr2*, and *Atp2a2* in ND-Mat and HFD-Mat offspring at P20 and P60 (N=10 rats/group). **(C)** Representative immunoblot and quantification of SERCA and JPH2 protein expression, normalized to total protein, at P20 and P60 in ND-Mat and HFD-Mat offspring (N=6 rats/group). **(D)** Representative immunofluorescence images of Cav3 (red) and Cav1.2 (green) and **(E)** of JPH2 (green) and Cav3 (red) in isolated cardiomyocytes from P60 ND-Mat and HFD-Mat offspring, with corresponding fluorescence intensity line-scan profiles illustrating periodicity of the fluorescence staining, and representative 3D-rendered images of Cav3 (red) and JPH2 (green) labeling (N=3 rats/group). **(F)** Representative immunofluorescence images of RyR2 (green), wheat germ agglutinin (WGA, red), and DAPI (blue) on transverse cardiac tissue sections from P60 ND-Mat and HFD-Mat offspring (N=3 rats/group). **(G)** Higher-magnification images and quantification of RyR2 labeling in longitudinal sections, expressed as the number of green pixels per cardiomyocyte area (N=3 rats/group). **(H)** Representative immunofluorescence images of Cav1.2 (green) and SERCA (red) in isolated cardiomyocytes from P60 ND-Mat and HFD-Mat offspring (N=3 rats/group). **(I)** Higher-magnification images and quantification of SERCA labeling, expressed as the number of green pixels per cardiomyocyte area (N=3 rats/group). Data are presented as mean±SD; two-way ANOVA followed by Tukey’s multiple comparisons test (B, C) and Unpaired Student’s *t*-test (G, I); \**P*<0.05; \*\**P*<0.01; \*\*\**P*<0.001; \*\*\**P*<0.0001; ns, not significant.

Western blot analyses confirmed these transcriptomic changes at the protein level, demonstrating increased expression of junctophilin-2 (JPH2) and SERCA2 at P20, followed by reduced expression of both proteins in adult HFD-Mat hearts at P60 (Figure 7C).

To determine whether the downregulation of Cav3 translated into structural alterations of the membrane, caveolin-3 immunofluorescence was performed on Langendorff-isolated cardiomyocytes. In P60 HFD-Mat cardiomyocytes, Cav3 labelling showed a loss of the regular, punctate pattern normally associated with caveolae and T-tubule invaginations, with an apparent flattening of the membrane ridges and T-tubule network along the cell, as confirmed by 3D reconstruction, suggestive of impaired caveolar and T-tubule organization (Figure 7D).

Because JPH2 is essential for maintaining the close apposition between T-tubules and the sarcoplasmic reticulum, its localization was also examined. Immunofluorescence revealed a marked loss of the regular striated JPH2 pattern in HFD-Mat cardiomyocytes, confirmed by fluorescence distribution analysis (Figure 7E), while 3D reconstruction showed a pronounced reduction of JPH2 signal in the central regions of the cell, consistent with disruption of the transverse tubular network (Figure 7E).

We next examined RyR2, the major sarcoplasmic reticulum Ca²⁺ release channel mediating calcium-induced calcium release during excitation-contraction coupling. In fresh cardiac biopsies from adult HFD-Mat hearts, RyR2 labelling was markedly decreased on transverse sections (Figure 7F), a reduction that was confirmed and quantified on longitudinal sections (Figure 7G).

In Langendorff-isolated cardiomyocytes, both the L-type calcium channel Cav1.2 and SERCA2a similarly lost their characteristic striated organization in HFD-Mat cells (Figure 7H). Consistent with this, SERCA2a labelling was also reduced in fresh cardiac biopsies on longitudinal sections (Figure 7I), with a similar reduction observed on transverse sections (Supplemental Figure 7C).

Together, these observations demonstrate that maternal HFD-Mat drives a profound remodeling of the excitation-contraction coupling machinery, providing a structural basis for the impaired systolic function observed in adult offspring.

### Maternal HFD-Mat promotes early intercalated disc maturation but disrupts adult intercalated disc organization (Figure 8)

Following the characterization of sarcomeric and E-C coupling apparatus remodeling, we next investigated the intercalated disc (ID), a specialized cell-cell junction that mechanically and electrically couples adjacent cardiomyocytes, thereby ensuring efficient force transmission and synchronous electrical propagation throughout the myocardium (Figure 8.A). Transcriptomic analysis revealed distinct, gene-specific regulatory dynamics among key ID components in HFD-Mat hearts: *Gja1* (Connexin-43) was significantly upregulated at P20 but returned to control levels by P60, while *Cdh2* (N-cadherin) was unchanged at P20 but became significantly upregulated at P60. In contrast, *Ctnnb1* (β-catenin) expression was significantly decreased at P20, followed by an upregulation at P60. (Figure 8B). At the protein level, Western blot analysis showed increased expression of Cx43 and N-cadherin in HFD-Mat offspring at the juvenile stage (P20), with no significant differences detected at P60, while PKP2 and β-catenin expression remained unchanged at both ages (Figure 8C).

**Figure 8:**
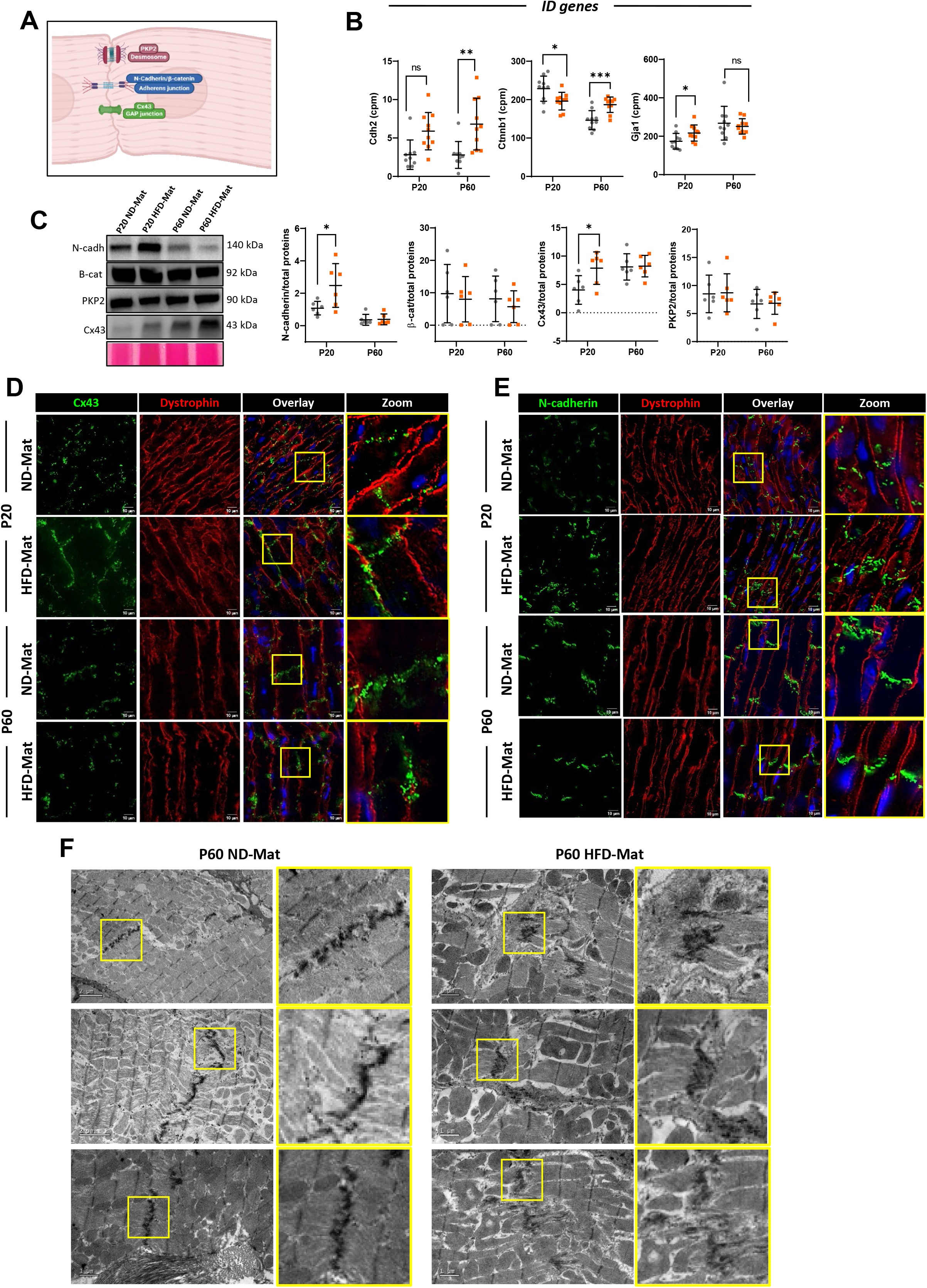
Maternal HFD promotes early intercalated disc maturation but disrupts adult intercalated disc organization. **(A)** Schematic representation of the main components of the intercalated disc. **(B)** Normalized RNA sequencing expression (counts per million, CPM) of genes encoding major intercalated disc proteins, including *Gja1* (connexin-43), *Cdh2* (N-cadherin) and *Ctnnb1* (β-catenin) at P20 and P60 stages (N=10 rats/group). **(C)** Representative immunoblots and corresponding quantification of the relative protein expression of Connexin-43 (Cx43), N-cadherin, β-catenin and PKP2 normalized to total protein staining (Ponceau S) (N=6 rats/group). **(D)** Representative immunofluorescence images of cardiac sections stained for Cx43 and **(E)** N-cadherin, showing their subcellular localization at P20 and P60 stages (N=6 rats/group). **(F)** Representative transmission electron microscopy images of the intercalated disc illustrating ultrastructural architecture in adult HFD-Mat offspring (N=3 rats/group). Data are presented as mean±SD; two-way ANOVA followed by Tukey’s multiple comparisons test (B, C); \**P*<0.05; \*\**P*<0.01; \*\*\**P*<0.001; \*\*\**P*<0.0001; ns, not significant.

To determine whether the absence of a protein-level difference at P60 reflected preserved ID architecture or masked an underlying reorganization, we examined the subcellular localization of these proteins by immunofluorescence.

At P20, Cx43 exhibited a more organized localization at the intercalated discs in HFD-Mat hearts, whereas ND-Mat myocardium displayed the less organized distribution expected at this immature developmental stage (Figure 8D). In contrast, despite comparable total Cx43 protein levels, P60 HFD-Mat hearts showed a marked lateralization of Cx43, with redistribution from the intercalated disc toward the lateral membrane of cardiomyocytes, whereas N-cadherin localization remained largely preserved (Figure 8E).

Ultrastructural analysis by transmission electron microscopy further confirmed disruption of ID architecture in adult HFD-Mat offspring (Figure 8F), supporting the progressive structural remodeling of cardiomyocyte cell-cell junctions.

Given that Cx43 lateralization has previously been linked to altered electrical conduction (Fontes et al. 2012), we next examined cardiac electrical function in adult offspring. Baseline electrocardiographic recordings at P60 revealed a significant increase in the RR interval, consistent with reduced heart rate in HFD-Mat offspring (Supplemental Figure 8). In contrast, β-adrenergic stimulation with isoproterenol elicited comparable responses in both groups and no significant differences were detected for the remaining ECG parameters (Supplemental Figure 8).

Together, these results show that maternal HFD-Mat accelerates intercalated disc maturation in early life but drives its progressive disorganization in adulthood, characterized by lateral redistribution of Cx43 despite unchanged total protein levels. Collectively, our results position the cardiomyocyte cytoarchitecture — sarcomere, excitation-contraction machinery and intercalated disc alike — as a coordinated target of maternal metabolic programming.

## DISCUSSION

The present study demonstrates that maternal HFD-Mat profoundly remodels postnatal cardiac development. Maternal HFD-Mat accelerates postnatal cardiac maturation, generating a transient adaptive phenotype that ultimately predisposes the heart to adult contractile dysfunction. Early after birth, offspring exposed to maternal HFD-Mat displayed coordinated maturation of metabolic pathways, sarcomere organization and excitation–contraction coupling, together with increased stroke volume and cardiac output and concentric physiological hypertrophy, accompanied by preserved or even enhanced systolic function at P20. However, these adaptations were not sustained and progressively evolved toward impaired cardiac performance in adulthood, characterized by reduced contractility, disorganization of the contractile apparatus, remodeling of the intercalated disc and lateral membrane, and alterations in mitochondrial homeostasis. Importantly, these defects occurred in the absence of overt fibrosis, myocardial lipid accumulation or major mitochondrial respiratory dysfunction previously reported to be triggered by exposure to HFD-Mat^20,21^, suggesting that disruption of cardiomyocyte architecture precedes the classical hallmarks of pathological remodeling. Because offspring were never directly exposed to HFD-Mat after weaning, these mechanisms likely differ from those triggered by direct HFD-Mat consumption, suggesting that maternal HFD-Mat instead engages specific developmental signals whose nature remains to be determined.

The transition from the fetal glycolytic heart toward the fatty acid-dependent adult myocardium is normally accompanied by extensive structural and metabolic maturation^22,23^. Our findings suggest that maternal metabolic stress shifts the timing of these developmental programs, promoting an earlier acquisition of adult-like metabolic and contractile features. This accelerated maturation may represent an adaptive response that enables the neonatal heart to cope with an altered postnatal nutritional environment. However, by shortening the normal developmental window, it may also reduce cardiac plasticity and increase susceptibility to dysfunction later in life. Consistent with this hypothesis, adult HFD-Mat offspring developed systolic dysfunction at P60 despite the absence of ventricular dilatation or overt diastolic impairment at this stage, with reduced intrinsic myocardial contractility indicating that functional deterioration primarily reflects impaired cardiomyocyte performance rather than secondary ventricular remodeling. Notably, because all offspring were weaned onto a standard diet irrespective of maternal feeding regime, this progressive trajectory unfolds entirely in the absence of any ongoing dietary exposure, indicating that the developmental window encompassing gestation and lactation is sufficient to permanently reprogram the postnatal maturation trajectory of the heart. These findings support the Developmental Origins of Health and Disease (DOHaD) hypothesis, according to which developmental adaptations promoting short-term survival may become maladaptive in adulthood when environmental demands change^24,25^. In this context, the early adaptive phenotype and the subsequent contractile dysfunction likely represent sequential stages of the same developmental trajectory rather than independent outcomes.

This developmental reprogramming of the heart occurred largely independently of any overt systemic metabolic disease we could measure. HFD-Mat offspring displayed normal glucose tolerance and insulin sensitivity, together with unchanged systolic blood pressure throughout postnatal life, despite the well-established epidemiological association between maternal obesity and offspring risk of diabetes and hypertension^3,26^. This dissociation suggests that maternal HFD-Mat programs the heart directly, rather than acting through systemic insulin resistance or hypertension, making cardiac cytoarchitecture an early, possibly independent, target of this programming.

One of the major findings of this study is the progressive remodeling of cardiomyocyte cytoarchitecture. Although sarcomere organization appeared enhanced at P20, adult HFD-Mat hearts exhibited marked sarcomeric disorganization together with reduced expression of several contractile genes. Interestingly, cardiomyocyte hypertrophy persisted despite this loss of structural organization, suggesting that cellular growth becomes dissociated from functional maturation during disease progression. Beyond this loss of overall organization, the coordination between neighboring sarcomeres was itself disrupted, with the physiological correlation in length between adjacent sarcomeres completely abolished in adult HFD-Mat cardiomyocytes — a defect in myofibrillar coherence that may constitute a structural substrate not only for impaired force transmission but also for arrhythmogenic vulnerability^27^.

This remodeling was not restricted to the contractile apparatus but also involved specialized membrane domains responsible for mechanical and electrical coupling. Connexin43, normally enriched at the intercalated disc, displayed abnormal lateral redistribution, indicating a loss of cardiomyocyte polarity. In parallel, reduced Caveolin-3 expression and the loss of membrane invaginations, together with decreased expression of key excitation–contraction coupling proteins including RyR2, SERCA2a and Junctophilin-2, demonstrate that early metabolic stress progressively disrupts the structural organization required for efficient calcium signaling, providing a structural basis for the contractile dysfunction of adult offspring. These alterations were paralleled by signs of altered cardiac electrical and autonomic regulation, including a divergent trajectory in heart rate variability in adult hearts, coincident with Cx43 lateralization. Because connexin distribution and autonomic tone jointly shape impulse conduction and chronotropy^28^, maternal HFD-Mat may program the heart’s electrical and autonomic control as well as its contractile machinery, consistent with reports of autonomic programming after early-life high-fat exposure in other rodent models^29^.

Given the established role of mitochondrial dysfunction in obesity-associated cardiomyopathy, we examined whether energetic failure contributed to the adult phenotype. Despite changes in mitochondrial morphology, reduced ATP content, increased Parkin Ser65 phosphorylation, and reduced CPT1B, overall mitochondrial respiratory capacity remained preserved. Fatty acid oxidation capacity was not directly assessed, so whether the moderate CPT1B reduction translates into functional impairment remains unknown and will require further investigation. Together, these findings point to activation of mitochondrial quality-control mechanisms rather than severe bioenergetic failure, indicating that mitochondrial dysfunction is unlikely to be the primary driver of contractile impairment in this model.

Diastolic function deteriorated later than systolic function, indicating that cardiac impairment continues to accumulate beyond the stages covered by most of our mechanistic analyses. This raises the possibility that the metabolic and structural alterations already detected at P60 — including mitochondrial morphology, ATP content and quality-control pathways — likewise represent early stages of a process that could progress further beyond P120. Long-term follow-up will be required to test this hypothesis.

Several limitations should be acknowledged. The molecular mechanisms underlying this developmental shift remain to be established; investigating the earliest postnatal stage (P5), and intermediate stages beyond P20 and P60, may help identify the initial events triggered by maternal HFD-Mat and clarify the temporal sequence of structural remodeling, potentially involving epigenetic, placental or endocrine mechanisms. Additionally, although transcriptomic, lipidomic and metabolomic analyses showed no sex-dependent clustering, our study was not powered to test for sex-by-diet interactions, and subtler sex-dimorphic effects on cardiac programming — reported in other developmental programming models ^30^— cannot be excluded.

Beyond mechanistic insights, our findings also carry potential clinical relevance. Maternal obesity affects an increasing proportion of pregnancies worldwide, yet cardiovascular surveillance of offspring remains limited. Plasma metabolomic and lipidomic analyses identified alterations in circulating fatty acids, including increased stearic and oleic acid, as early as P5 — well before the onset of overt cardiac dysfunction and at a stage approximately corresponding to infancy in humans. These circulating metabolites could represent early, minimally invasive candidate biomarkers to help identify offspring of mothers with overweight or obesity who may be at increased cardiovascular risk, an avenue that will require validation in future studies. Circulating ANP and BNP, already elevated by P60 alongside the onset of contractile dysfunction, could complement such a biomarker panel at later stages, consistent with their established use in human heart failure^31^.

In conclusion, our study demonstrates that maternal HFD-Mat reprograms postnatal cardiac development by accelerating, rather than delaying, cardiac maturation. This premature developmental transition is initially associated with adaptive structural and functional changes but ultimately predisposes the heart to adult systolic dysfunction through progressive remodeling of cardiomyocyte cytoarchitecture. Notably, our findings indicate that altered developmental timing, rather than primary metabolic failure, is the main feature associated with cardiac dysfunction in this model. By identifying early postnatal life as the period when these maladaptive trajectories begin, this work provides new insight into the developmental origins of cardiovascular disease and highlights the potential value of early diagnostic strategies for offspring exposed to maternal obesity.

## Supplemental Methods

### Maternal obesity accelerates cardiomyocyte maturation and programs adult heart dysfunction in offspring

Farhi et al. – Data Supplement

#### Animal model and study design

All rats used in this study, both adult males and females, were of the Wistar strain (Janvier Labs, France). Animals were housed under specific pathogen-free (SPF) conditions with ad libitum access to food and water, in a temperature-controlled room on a 12:12 h light/dark cycle. Female rats (mothers) were fed either a normal diet (ND-Mat group) or a high-fat diet (HFD-Mat group; D12492, Research Diets, Inc., USA) for two months prior to mating and throughout gestation and lactation. After weaning, all offspring were fed a standard chow diet regardless of maternal diet. Experimental studies were conducted at different postnatal stages (P5, P20, P60 and P120). All procedures were carried out in accordance with institutional guidelines (Ministère de l’Agriculture, France, authorization no. 75-1090) and the European Directive 2010/63/EU on the protection of animals used for scientific purposes.

#### Echocardiographic assessment of cardiac function

Echocardiographic analyses were performed on rats anesthetized with isoflurane inhalation (3–4% isoflurane in 1 L/min oxygen). Cardiac function was further challenged by an intraperitoneal injection of isoproterenol (50 mg/kg; Sigma-Aldrich, I5627-25G) diluted in 0.9% NaCl solution. Image acquisition and measurements were performed using a Vevo 3100 ultrasound imaging system (VisualSonics) at the UMS28 core facility (Pitié-Salpêtrière Hospital, Paris, France). M-mode, Doppler, and tissue Doppler imaging modalities were used for this study.

#### Electrocardiogram assessment

Electrocardiograms (ECGs) were recorded in rats anesthetized with isoflurane inhalation (3–4% isoflurane in 1 L/min oxygen) at the three postnatal stages using needle electrodes (ecgAIGUILLES, Emka Technologies). Recordings were acquired with the IOX 2.10.040 software (Emka Technologies). The duration of the P wave and the PR, QRS, QT, and RR intervals was automatically measured using the ECG Auto software (ECGAUTO v3.0, Emka Technologies). To assess the chronotropic response, a challenge with isoproterenol (50 mg/kg; I5627-25G, Sigma-Aldrich) diluted in 0.9% NaCl was performed by intraperitoneal injection. ECG recordings were continued for 4 minutes until the plateau phase was reached and heart rate returned to baseline levels. RR intervals were also analyzed using Poincaré plots generated with a custom script in R. Each RR interval (RRn) was plotted against the subsequent interval (RRn+1), providing a graphical assessment of beat-to-beat heart rate variability. The standard descriptors SD1 and SD2 were calculated to quantify short-term and long-term RR interval variability, respectively.

#### Invasive hemodynamic measurements

P60 stage offspring rats were anesthetized with a mix of ketamine/xylazine (100 mg/kg/10 mg/kg, IP). Animals were placed on controlled heating pads, and the temperature was maintained at 37°C. Invasive hemodynamic measurements were then performed on spontaneously breathing rats using a pressure transducer catheter (SPR-838, 2F Millar Mikro-tip P-V (pressure-volume) catheter, Millar Instruments) introduced into the right carotid artery and advanced into the LV (left ventricle) under pressure control. Signals were continuously recorded at a sampling rate of 1000 samples/s using a P-V conductance system (MVPS-Ultra, Millar Instruments) connected to a PowerLab 16/30 data acquisition system (AD Instruments), stored, and displayed by the Lab-Chart 5 Software System (AD Instruments). LV P-V relations were also registered during transient compression of the inferior vena cava (reducing preload) using a syringe pusher. Indices of LV contractility (ESPVR and PRSW) and compliance (EDPVR) were analysed using P-V analysis program (PVAN; Millar Instruments).

#### Isolation and culture of rat cardiomyocytes

Adult ventricular cardiomyocytes were isolated from 3-weeks-old (P20) and 2-month-old (P60) Wistar rats using Langendorff retrograde perfusion. Following heparinization (250 U/kg, i.p.) and euthanasia with Euthasol (120 mg/kg), hearts were rapidly excised, cannulated via the ascending aorta, and perfused at 37°C. Coronary circulation was first flushed with oxygenated Krebs–Henseleit buffer supplemented with 15 mM 2,3-butanedione monoxime (BDM) and 10 mM taurine, followed by enzymatic digestion with collagenase A (200 U/mL) in Krebs–Henseleit buffer containing 15 mM BDM and 7.5 mg/mL bovine serum albumin (BSA). Ventricles were mechanically dissociated in Kraft–Brühe (KB) buffer to obtain isolated cardiomyocytes.

For high-resolution respirometry, mitochondrial membrane potential measurements, and cell death assays, cells were plated on poly-D-lysine- and laminin-coated coverslips or culture dishes and maintained at 37°C in 5% CO₂. After 90 min, KB buffer was replaced with M199 medium supplemented with 1% fetal bovine serum (FBS), 1% penicillin/streptomycin, and 1% insulin–transferrin–selenium (ITS). Cardiomyocytes were subsequently transferred to the Mitochondrial Biology Unit, Institut Pasteur (UMR 3691, Paris, France), for mitochondrial functional analyses.

#### Glucose and insulin tolerance tests

For the intraperitoneal glucose tolerance test (IPGTT), rats were fasted for 6 hours before receiving an intraperitoneal injection of D-glucose (2 g/kg; Sigma-Aldrich, G7528) diluted in sterile saline. For the intraperitoneal insulin tolerance test (IPITT), rats were also fasted for 6 hours prior to the experiment and received an intraperitoneal injection of insulin (0.75 U/kg; Actrapid, Novo Nordisk). For both tests, blood samples were collected from the tail vein at 0, 15, 30, 60, 90, and 120 minutes after injection, and blood glucose concentrations were measured using a glucometer (Accu-Chek Performa, Roche).

#### Plasma biochemical analyses

Venous blood was collected by right ventricular puncture immediately after euthanasia using heparinized syringes (26G × 12 mm, Terumo). Samples were centrifuged at 3,000 × g for 15 min at 4°C, and plasma was collected, aliquoted, and stored at −80°C until analysis.

Plasma concentrations of glucose, triglycerides, total cholesterol, and C-reactive protein (CRP) were measured using an Indiko™ automated clinical chemistry analyzer (Thermo Fisher Scientific, Waltham, MA, USA) according to the manufacturer’s instructions with the corresponding commercial reagent kits (Thermo Fisher Scientific). Quality control procedures and instrument calibration were performed following the manufacturer’s recommendations.

#### Plasma ANP and BNP measurements

Plasma concentrations of atrial natriuretic peptide (ANP) and B-type natriuretic peptide (BNP) were quantified at P20 and P60 using commercially available ELISA kits (ANP, RAB0385; BNP, RAB0386-1KT; Sigma-Aldrich, St. Louis, MO, USA) according to the manufacturer’s instructions. Plasma samples were assayed in duplicate, and absorbance was measured using a microplate reader. ANP and BNP concentrations were calculated from standard curves generated using the standards supplied with each kit.

#### Cardiac ATP quantification

Cardiac ATP content was quantified in P60 heart tissue using the ATP Determination Kit (A22066, Thermo Fisher Scientific, Waltham, MA, USA) according to the manufacturer’s instructions. Briefly, approximately 100 mg of cardiac tissue was homogenized in ultrapure water using bead-based homogenization (2 min, 50 Hz). Tissue lysates were heated at 95°C for 5 min to inactivate ATP-degrading enzymes before analysis. Bioluminescence was measured using a Tecan microplate reader, and ATP concentrations were determined from a standard curve generated with ATP standards provided in the kit.

#### Blood pressure measurement

Systolic blood pressure was assessed non-invasively by tail-cuff plethysmography using the BP-2000 Blood Pressure Analysis System (Bioseb). Measurements were performed at P20 and P60 after a 4-day acclimation period to minimize stress-related effects.

#### Protein extraction and Western Blot Analysis

Rats were euthanized through an intraperitoneal injection of Euthasol (140mg/kg). Frozen myocardial tissues were homogenized in the following lysis buffer (in mmol/L): 150 NaCl, 50 Tris-HCl (pH 7.5), 1 EDTA, 0.5% Triton X-100, and 1% protease inhibitor cocktail (Sigma-Aldrich). Solubilization was performed by rotating the samples for 2 hours at 4 °C. Supernatants obtained after centrifugation at 13,000 g for 15 min at 4 °C were collected for further analysis. Protein concentrations were determined using the BCA Protein Assay Kit (23227, Thermo Fisher Scientific) with a BSA standard curve. After extraction, proteins were separated on NuPAGE® Novex® 10% Bis-Tris gels (Life Technologies) and transferred onto nitrocellulose membranes using the iBlot® 2 Dry Blotting System. Membranes were incubated with the appropriate primary antibodies (cf. Table I), followed by HRP-conjugated secondary antibodies (Cell Signaling Technology). Protein bands were visualized using ECL Prime detection reagent (GE Healthcare), and images were acquired with a LAS4000 imaging system (GE Healthcare). Since no stable loading control protein was identified between the ND and HFD-Mat groups, target protein levels were normalized to total protein content using Ponceau Red staining (P7170, Sigma-Aldrich). For OxPhos protein detection, membranes were heated for 10 minutes at 37 °C prior to incubation with a mouse anti-OxPhos antibody cocktail (1:1000; Invitrogen, Cat# 45-8099, RRID: AB_2533835).

**Table 1:** List of antibodies.

| Target protein | Antibody type, host specie | Reference |
| --- | --- | --- |
| **α-actinine** | Mouse monoclonal | 7811 (Abcam) |
| **β-caténine** | Mouse monoclonal | Ab22656 (Abcam) |
| **Cav1.2** | Rabbit polyclonal | ACC-003 (Alomone) |
| **Caveolin-3** | Mouse monoclonal | 610420 (BD Transduction Lab) |
| **Connexin 43** | Rabbit polyclonal | 710700 (Lifetech) |
| **CPT1B** | Rabbit polyclonal | 22170-1-AP (Prointech) |
| **Desmin** | Mouse monoclonal | D-1033 (Sigma-Aldrich) |
| **Dystrophin** | Mouse monoclonal | Ab3149 (Abcam) |
| **Dystrophin** | Mouse monoclonal | Ab15277 (Abcam) |
| **Hexokinase 1** | Rabbit polyclonal | Ab227198 (Abcam) |
| **JPH2** | Rabbit polyclonal | 40-5300 (Thermo) |
| **N-cadherin** | Mouse monoclonal | 33-3900 (Thermo) |
| **OXPHOS** | Mouse monoclonal | 45-8099 (Invitrogen) |
| **Parkin-S65** | Rabbit monoclonal | Ab315376 (Abcam) |
| **Ryr2** | Mouse monoclonal | Ab2827 (Abcam) |
| **SERCA2 ATPase** | Mouse monoclonal | MA3-919 (Invitrogen) |
| **Cardiac troponine T** | Rabbit monoclonal | Ab45932 (Abcam) |

**Table 2.**
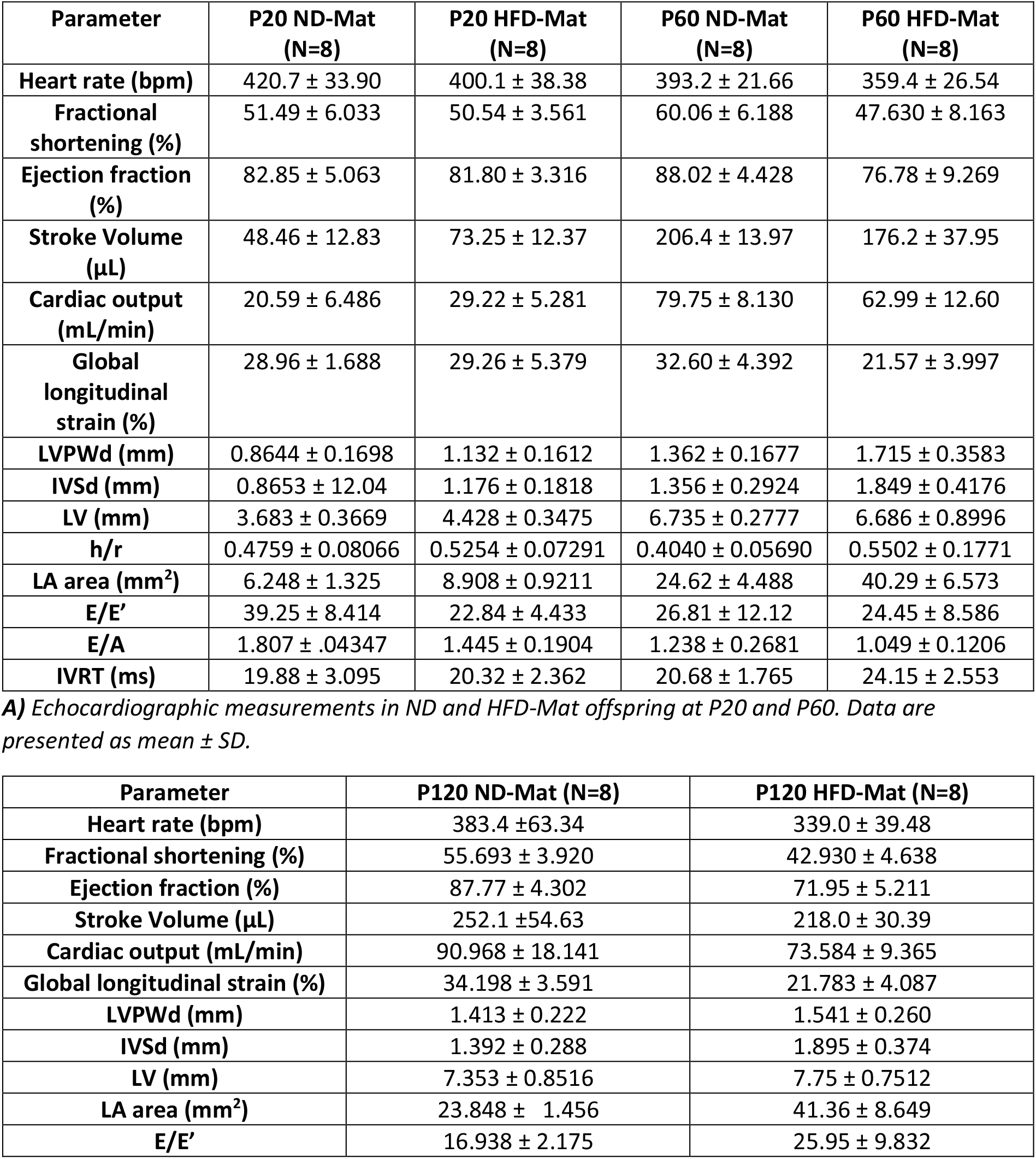
Echocardiographic parameters.

| Parameter | P20 ND-Mat (N=8) | P20 HFD-Mat (N=8) | P60 ND-Mat (N=8) | P60 HFD-Mat (N=8) |
| --- | --- | --- | --- | --- |
| Heart rate (bpm) | 420.7 ± 33.90 | 400.1 ± 38.38 | 393.2 ± 21.66 | 359.4 ± 26.54 |
| Fractional shortening (%) | 51.49 ± 6.033 | 50.54 ± 3.561 | 60.06 ± 6.188 | 47.630 ± 8.163 |
| Ejection fraction (%) | 82.85 ± 5.063 | 81.80 ± 3.316 | 88.02 ± 4.428 | 76.78 ± 9.269 |
| Stroke Volume (μL) | 48.46 ± 12.83 | 73.25 ± 12.37 | 206.4 ± 13.97 | 176.2 ± 37.95 |
| Cardiac output (mL/min) | 20.59 ± 6.486 | 29.22 ± 5.281 | 79.75 ± 8.130 | 62.99 ± 12.60 |
| Global longitudinal strain (%) | 28.96 ± 1.688 | 29.26 ± 5.379 | 32.60 ± 4.392 | 21.57 ± 3.997 |
| LVPWd (mm) | 0.8644 ± 0.1698 | 1.132 ± 0.1612 | 1.362 ± 0.1677 | 1.715 ± 0.3583 |
| IVSd (mm) | 0.8653 ± 12.04 | 1.176 ± 0.1818 | 1.356 ± 0.2924 | 1.849 ± 0.4176 |
| LV (mm) | 3.683 ± 0.3669 | 4.428 ± 0.3475 | 6.735 ± 0.2777 | 6.686 ± 0.8996 |
| h/r | 0.4759 ± 0.08066 | 0.5254 ± 0.07291 | 0.4040 ± 0.05690 | 0.5502 ± 0.1771 |
| LA area (mm<sup>2</sup>) | 6.248 ± 1.325 | 8.908 ± 0.9211 | 24.62 ± 4.488 | 40.29 ± 6.573 |
| E/E' | 39.25 ± 8.414 | 22.84 ± 4.433 | 26.81 ± 12.12 | 24.45 ± 8.586 |
| E/A | 1.807 ± .04347 | 1.445 ± 0.1904 | 1.238 ± 0.2681 | 1.049 ± 0.1206 |
| IVRT (ms) | 19.88 ± 3.095 | 20.32 ± 2.362 | 20.68 ± 1.765 | 24.15 ± 2.553 |

**A)** Echocardiographic measurements in ND and HFD-Mat offspring at P20 and P60. Data are presented as mean ± SD.
| Parameter | P120 ND-Mat (N=8) | P120 HFD-Mat (N=8) |
| --- | --- | --- |
| Heart rate (bpm) | 383.4 ± 63.34 | 339.0 ± 39.48 |
| Fractional shortening (%) | 55.693 ± 3.920 | 42.930 ± 4.638 |
| Ejection fraction (%) | 87.77 ± 4.302 | 71.95 ± 5.211 |
| Stroke Volume (μL) | 252.1 ± 54.63 | 218.0 ± 30.39 |
| Cardiac output (mL/min) | 90.968 ± 18.141 | 73.584 ± 9.365 |
| Global longitudinal strain (%) | 34.198 ± 3.591 | 21.783 ± 4.087 |
| LVPWd (mm) | 1.413 ± 0.222 | 1.541 ± 0.260 |
| IVSd (mm) | 1.392 ± 0.288 | 1.895 ± 0.374 |
| LV (mm) | 7.353 ± 0.8516 | 7.75 ± 0.7512 |
| LA area (mm<sup>2</sup>) | 23.848 ± 1.456 | 41.36 ± 8.649 |
| E/E' | 16.938 ± 2.175 | 25.95 ± 9.832 |

#### Immunofluorescence

Myocardial cryosections (7 µm) were incubated for 1 hour in a permeabilization/blocking buffer containing PBS, 0.5% Triton X-100, 1% BSA, and 10% serum from the species in which the secondary antibodies were raised. Sections were then incubated overnight at 4 °C with primary antibodies (cf. Table I) diluted in permeabilization/blocking buffer (PBS, 0.5% Triton X-100, 1% BSA, 3% serum). After three washes in PBS, sections were incubated for 1 hour at room temperature, protected from light, in PBS containing 1% BSA and 3% serum, together with secondary antibodies conjugated to Alexa Fluor 488, 546, or 594 (2.5–5 µg/mL; Jackson ImmunoResearch), and the nuclear and membrane markers DAPI (0.2 µg/mL; Sigma-Aldrich) and WGA (1:400; Sigma-Aldrich), respectively. Mounting was performed using Dako Fluorescence Mounting Medium.

#### Fluorescent immunolabeling and confocal imaging of fresh cardiac biopsies

Fresh cardiac biopsies were immunolabeled and imaged using the 3D-NaissI (3D-Native Tissue Imaging) method, as previously described^17^. Briefly, fresh biopsies were incubated in HBSS with the primary antibody for 2 days at 4 °C under constant rotary agitation, then washed and incubated with the fluorescent secondary antibody, protected from light, for a further 2 days under the same conditions before imaging.

Biopsies were imaged on an inverted ZEISS LSM 900 Axio Observer.Z1/7 confocal microscope using a ×63 oil-immersion objective (NA 1.4) at a maximum resolution of 1437 × 1437 pixels (pixel scaling 0.071 × 0.071 µm). Three-dimensional stacks (10 µm thickness) were acquired with a z-step of 1 µm and an x/y pixel size of 0.099 µm. Emission filter ranges, detector gains, and pixel dwell time (0.54 µs) were set as detailed in^17^. Images were visualized with Zen Blue software (Carl Zeiss, Germany). For comparative studies, acquisition parameters were kept identical for a given primary antibody, but could differ between antibodies. Images presented correspond to the most representative selection among several independent acquisitions.

For each biopsy, three 2D image stacks were selected (top, middle, and bottom of the sample), each capturing the same cardiomyocytes (CMs) (i.e., three stacks per CM). Two-dimensional fluorescence images (transverse sections of left ventricular myocardium, CMs only) were analyzed with a custom Fiji/ImageJ macro enabling automated cell-by-cell quantification. After import via Bio-Formats, CMs were segmented with Cellpose based on fluorescent-wheat germ agglutinin (WGA) surface staining (CM mask), generating a label image converted to 16-bit for subsequent analysis.

To quantify total expression of the protein of interest in each CM, a binary segmentation of the protein fluorescence signal was obtained by defining a threshold in the control condition for each individual experiment. Protein expression was quantified as the number of fluorescent pixels above threshold rather than as mean fluorescence intensity. This pixel-based approach was chosen because the tissue and inter-experiment variability inherent to the 3D-NaissI method precludes reliable comparison of absolute fluorescence intensities^17^.

Accordingly, for each CM, the number of fluorescent pixels corresponding to the protein of interest was measured by 2D region analysis (MorphoLibJ) across three independent stacks per biopsy (5 biopsies per rat; 3 stacks per biopsy; ∼50–100 CMs per rat), and results were exported in CSV format. The total amount of fluorescent protein for a given condition was calculated as the total number of fluorescent pixels per CM per stack, summed across the three independent stacks per biopsy.

#### Sirius Red staining

Tissues were fixed in 4% paraformaldehyde (PFA; Sigma-Aldrich) for 48 hours under gentle rotation at 4 °C, followed by two successive washes in 70% ethanol, two washes in 90% ethanol, and three washes in absolute ethanol (each wash lasting 20 minutes). After dehydration, hearts were cleared in methyl salicylate, with a first incubation for 2 hours and a second overnight. Hearts were then embedded in paraffin (Leica). Histological staining was performed on 7 µm sections obtained using a microtome. Paraffin-embedded myocardial sections were incubated for 1 hour at room temperature in Sirius Red solution (365548-25G, Sigma-Aldrich), followed by two washes in 0.01% hydrochloric acid. Sections were then dehydrated in 100% ethanol and xylene, and mounted using Eukitt mounting medium (03989, Sigma-Aldrich).

#### Oil Red O staining

Myocardial cryosections were incubated in 60% 2-propanol for 20 seconds, followed by 10 minutes in Oil Red O solution (1024190250, Sigma-Aldrich). Sections were then briefly incubated again in 60% 2-propanol for 30 seconds and rinsed in distilled water for 20 seconds. Haematoxylin counterstaining (GHS316, Sigma-Aldrich) was performed by incubating sections for 5 minutes. Finally, sections were rinsed under running water for 3 minutes and mounted with an aqueous mounting medium (F4680, Sigma-Aldrich).

#### Microscopy and image analysis

##### Deconvolution Microscopy

Myocardial cryosections were examined using a DeltaVision™ Elite epifluorescence microscope (GE Healthcare) equipped with a CoolSnap HQ2 monochrome EMCCD camera, a 40x and 60x/1.42 oil immersion Plan APO objective, and the softWoRx software for rapid deconvolution to acquire Z-stack digital images at 0.1 µm intervals. Images were processed and analysed using ImageJ software (NIH). For image representation, three deconvolved Z-slices were projected into a single composite image. Longitudinal myocardial regions were selected to localize proteins of interest within cardiomyocytes, while transverse regions were used to quantify cell size.

Deconvoluted image stacks were visualized in three dimensions using Imaris (Bitplane/Oxford Instruments, v10.1.0). The Surface module was used to render the fluorescence signal of interest, generating a texturized 3D surface reconstruction from the raw voxel intensities. The resulting 3D projection was rotated to display the labelling from an alternative viewing angle.

#### Optical Microscopy

Histological images were acquired using a Nikon Eclipse Ti optical microscope equipped with a 10x objective and controlled by the NIS-Elements software (Nikon Instruments, Tokyo, Japan). Fibrosis analysis was performed using HistoLab software by manually delineating the myocardial area. Automatic thresholding was then applied to quantify fibrotic and myocardial surfaces.

#### Electron microscopy

Samples were postfixed in 1% osmium tetroxide 0.1M (Electron Microscopy Science, UK) in 0.1 M Phosphate Buffer (PB) (pH 7.4). Samples were washed 3 times in H_2_O then dehydrated in alcohol grades: 70% Ethanol 10 min, 90% Ethanol 10 min, 100% Ethanol 3×15 min, 100% Propylene oxide (Electron Microscopy Science, UK) 2X5min. Resin infiltration was performed as following: mix 1:1 Epikote 812: propylene oxide 30 min followed by mix 1:2 Epikote 812: propylene oxide overnight at room temperature. Samples were washed in 100% Epikote 812 then embedded in silicon molds in 100% Epikote 812, and polymerised in 60°C oven for 24 hours. Ultrathin sections were cut at 90 nm with a Leica UFC7 ultramicrotome (Leica Microsystems GmbH, Germany) and deposed on Gilder grids 200 mesh (Electron Microscopy Science, UK). They were counterstained with uranyl acetate 7% (LFG, France) and Reynold’s lead citrate (LFG, France). Samples were examined in a JEOL 1011 transmission electron microscope (JEOL, Japan) with an ORIUS 1000 CCD camera (GATAN, France), operated with Digital Micrograph software (GATAN, France) for acquisition. Mitochondrial morphology was quantified in Fiji (ImageJ). Area, circularity, aspect ratio and solidity were measured on 45 to 50 images per group and per stage, taken at ×5,000 magnification.

#### High-resolution respirometry

Adult ventricular cardiomyocytes were isolated and cultured as described above. Mitochondrial respiration was assessed in freshly isolated adult cardiomyocytes using an O2k high-resolution respirometer (Oroboros Instruments, Innsbruck, Austria). Briefly, 20,000 cardiomyocytes were resuspended in Mir05 buffer [MgCl_2_-6H_2_O 3 mM, Lactobionic Acid 60 mM, Taurine 20 mM, KH_2_PO_4_ 10 mM, HEPES-KOH 20 mM, Sucrose 110 mM, EGTA-KOH 0.5 mM, BSA (1g/L)] and permeabilized with 1 µg of digitonin. Maximal mitochondrial respiration capacity (OXPHOS) was determined under consecutive administrations of PGM [pyruvate 10 mM, glutamate 5 mM, malate 5 mM, state 2] or malate (2 mM) and palmitoyl-carnitine (10 µM) in presence of ADP (1 mM, state 3) to assess complex I-driven respiration; rotenone (0.5 µM) and succinate (10 mM, state 2) for the measurement of complex II-driven respiration; antimycin A (2.5 µM), ascorbate (2 mM), *N*,*N*,*N*′,*N*′-tetramethyl-*p*-phenylenediamine (TMPD, 0.5 mM for the determination of the complex IV-driven respiration). Data were analysed using Datlab v.7.

#### Cell death assay

Freshly isolated cardiomyocytes were stained with NucBlue™ Live ReadyProbes™ (Thermo Fisher Scientific) and tetramethylrhodamine ethyl ester (TMRE, 50 nM) for 30 min at 37°C. Live-cell imaging was performed in the presence or absence of hydrogen peroxide (10 µM) or doxorubicin (60 µM) using an Operetta CLS high-content imaging system (PerkinElmer). Images were acquired hourly for 15 h using a 5x air objective. Cardiomyocyte viability was quantified using a supervised machine-learning algorithm implemented in Harmony software (v4.9, PerkinElmer) based on TMRE fluorescence intensity, cell morphology, and nuclear number.

#### Mitochondrial membrane potential

Mitochondrial membrane potential (ΔΨm) was evaluated in isolated cardiomyocytes by live-cell confocal imaging. Cells were plated onto laminin-coated CellCarrier Ultra 96-well plates (PerkinElmer), stained with NucBlue™ Live ReadyProbes™, TMRE (100 nM), and MitoTracker™ Deep Red (100 nM) for 30 min at 37°C, and imaged using an Operetta CLS microscope (PerkinElmer) equipped with 5x or 40x air objectives. TMRE and MitoTracker Deep Red fluorescence intensities were quantified on a per-cell basis using Harmony software (v4.9, PerkinElmer), and mitochondrial membrane potential was expressed as the TMRE/MitoTracker Deep Red fluorescence ratio.

#### RNA sequencing and bioinformatic analysis

Total RNA was extracted from the hearts of Wistar rats at P5, P20, and P60 stages born to mothers fed either a normal diet (ND-Mat) or a high-fat diet (HFD-Mat) (n = 10 per group and stage; 5 males and 5 females). Before euthanasia, animals received an intraperitoneal injection of heparin sodium (250 U/kg; H3149, Sigma-Aldrich) and were euthanized with Euthasol (120 mg/kg). Hearts were rapidly excised, snap-frozen, and stored at −80 °C until RNA extraction. Total RNA was isolated using TRIzol reagent (15596026, Thermo Fisher Scientific) according to the manufacturer’s instructions. mRNA libraries were prepared using the Illumina Stranded mRNA Library Prep Kit, and library quality was assessed using an Agilent 4200 TapeStation. Paired-end sequencing (2 × 150 bp) was performed on a NovaSeq X Plus platform (Illumina), generating an average of 40 million reads per sample after demultiplexing.

Raw sequencing reads were quality-checked using FastQC and processed with fastp for adapter trimming and quality filtering. Reads were aligned to the Rattus norvegicus rn7 reference genome using the Illumina DRAGEN Bio-IT Platform (v3.10.4), and gene counts were generated based on the GENCODE mRatBN7.2 annotation. Differential gene expression analysis was performed in R using DESeq2 (v1.26.0). Genes with low expression were filtered prior to normalization, and differential expression was assessed using a generalized linear model. Differentially expressed genes were defined by an absolute log₂ fold change > 1 and a Benjamini–Hochberg false discovery rate (FDR) < 0.05. Gene set enrichment analysis (GSEA) was performed using clusterProfiler (v3.14.3) and the KEGG pathway database.

#### Metabolomic analysis

Untargeted metabolomic profiling was performed on heart tissue and plasma collected from P5, P20, and P60 offspring born to normal diet (ND-Mat) or high-fat diet (HFD-Mat) mothers (n = 6 per group and stage; 3 males and 3 females). Cardiac tissues were rapidly excised, snap-frozen in liquid nitrogen, and stored at −80 °C. Plasma samples were isolated by centrifugation and stored at −80°C until analysis. Metabolites were extracted using ice-cold acetonitrile containing stable isotope-labelled internal standards. Analyses were performed by liquid chromatography–high-resolution mass spectrometry (LC-HRMS) using an Acquity UPLC system (Waters, Milford, MA, USA) coupled to a Q Exactive mass spectrometer (Thermo Fisher Scientific, Bremen, Germany). Chromatographic separation was achieved on a SeQuant ZIC-pHILIC column (Merck). Data were quality-controlled using pooled quality control samples, normalized, and processed prior to statistical analysis. Metabolite annotation was performed using reference databases including HMDB and KEGG.

#### Lipidomic analysis

Free fatty acid profiling was performed on heart tissue and plasma collected from P5, P20, and P60 offspring born to normal diet (ND-Mat) or high-fat diet (HFD-Mat) mothers (n = 6 per group and stage; 3 males and 3 females). Cardiac tissues were rapidly excised, snap-frozen in liquid nitrogen, and stored at −80 °C, while plasma samples were isolated by centrifugation and stored at −80°C until analysis. Free fatty acids were extracted from tissue homogenates and plasma in the presence of deuterated internal standards, derivatized with pentafluorobenzyl bromide (PFBBr), and analysed by gas chromatography–mass spectrometry (GC–MS) using a FAST BPX70 capillary column coupled to a single quadrupole mass spectrometer operating in chemical ionization mode with methane.

#### Statistics

Statistical analyses were performed with GraphPad Prism (version 11, GraphPad Software). Data are presented as mean ± SD unless otherwise stated. Two groups were compared with an unpaired two-tailed Student’s t-test. For more than two groups, one-way or two-way ANOVA was followed by Tukey’s post-hoc test; three-way ANOVA (Supplemental Figures 6 and 8) and Pearson’s correlation were used where indicated in the legends. Normality of the data was checked with the Shapiro-Wilk test before each statistical test was applied. No outliers were identified, so no data were excluded. The unit of analysis is stated in each figure legend. Differences were considered significant at P<0.05 (*P<0.05, **P<0.01, ***P<0.001, ****P<0.0001).

## ACKNOWLEDGEMENTS

This work was supported by the Fondation pour la Recherche Médicale (FRM), which funded the PhD fellowship of Dounia Farhi (Bourse Jeanne-Philippe Béziat, grant number [PBR202306017902]), and by the Fédération Française de Cardiologie (FFC), through thematic funding awarded to E.B. for this project (2024–2026). The authors also thank the Philanthropy Department of the IHU ICAN (Institute of Cardiometabolism and Nutrition), and in particular Francine Trocmé, for securing complementary funding for this work.

The authors thank Marie Lagouge for her expert guidance on mitochondrial biochemistry and for directing us toward Oroboros respirometry, Team 5, UMRS1166; Maria Mihoc and Olivier Bregerie for their assistance with animal husbandry and care; Maryse Guérin and Sophie Galier, Team 5, UMRS1166, for their help with the INDIKO experiments; Fabrice Atassi for his assistance with tail-cuff plethysmography, Team 3, UMRS1166; and Nadine Suffee, Raoul Manuel, Stéphane Hatem and Sophie Nadaud, Team 3, UMRS1166, for helpful discussions. The authors also thank Alain Schmitt for electron microscopy, Institut Cochin, INSERM U1016, CNRS UMR 8104; Farid Ichou and Marie L’Homme for lipidomics and metabolomics analyses, and Maharajah Ponnaiah for the biostatistical analysis of these data, ICAN Omics – Metabolomics platform and ICAN I/O – Data Sciences platform, Institute of Cardiometabolism and Nutrition (IHU ICAN); and the iGenSeq core facility, Institut du Cerveau (ICM, Paris, France), for genotyping and sequencing services, with particular thanks to Yannick Marie. Part of this work was carried out at the Data Analysis Core (RRID:SCR_026138). We gratefully acknowledge Marieke Vromman for assistance with RNAseq data analysis.

## AUTHORS CONTRIBUTION

D.F. performed all experiments, analyzed the data, and wrote the manuscript. G.D. performed experiments and analyzed some of the results. E.D. performed Oroboros and Operetta (cell death) experiments, analyzed the corresponding results, and wrote the associated Methods sections; she also reviewed and critically revised the manuscript. N.P. performed imaging on fresh biopsies and analyzed the data. L.M. developed the macro and performed the Poincaré plot analysis. C.B. performed preliminary experiments on the project and reviewed the manuscript. N.M. performed the echocardiography, pressure-volume loop, and ECG experiments in animals. T.W. provided expertise for the Oroboros and Operetta experiments, critically reviewed the manuscript, and provided funding for these aspects of the study. C.G. provided expertise in fresh-biopsy microscopy and theoretical input for project development. E.B. supervised the project, performed some experiments, wrote the manuscript, and obtained funding.

## COMPETING INTERESTS

The authors declare no competing interests.

## DATA AVAILABILITY

The RNA sequencing data generated in this study have been deposited in the Gene Expression Omnibus (GEO) database under accession number [GSE348573]. The lipidomic data generated in this study have been deposited in the Metabolomics Workbench repository (https://www.metabolomicsworkbench.org/data/index.php) under Project ID [PR003387, Study ID: ST005249, doi: 10.21228/M8WP17]; and the metabolomic data under Data Track ID 8155. All other data supporting the findings of this study are available within the article and its Supplementary Information, or from the corresponding author upon reasonable request. Source data are provided with this paper.

## DATA SUPPLEMENT – SUPPLEMENTAL FIGURE LEGENDS

**Figure 1:**
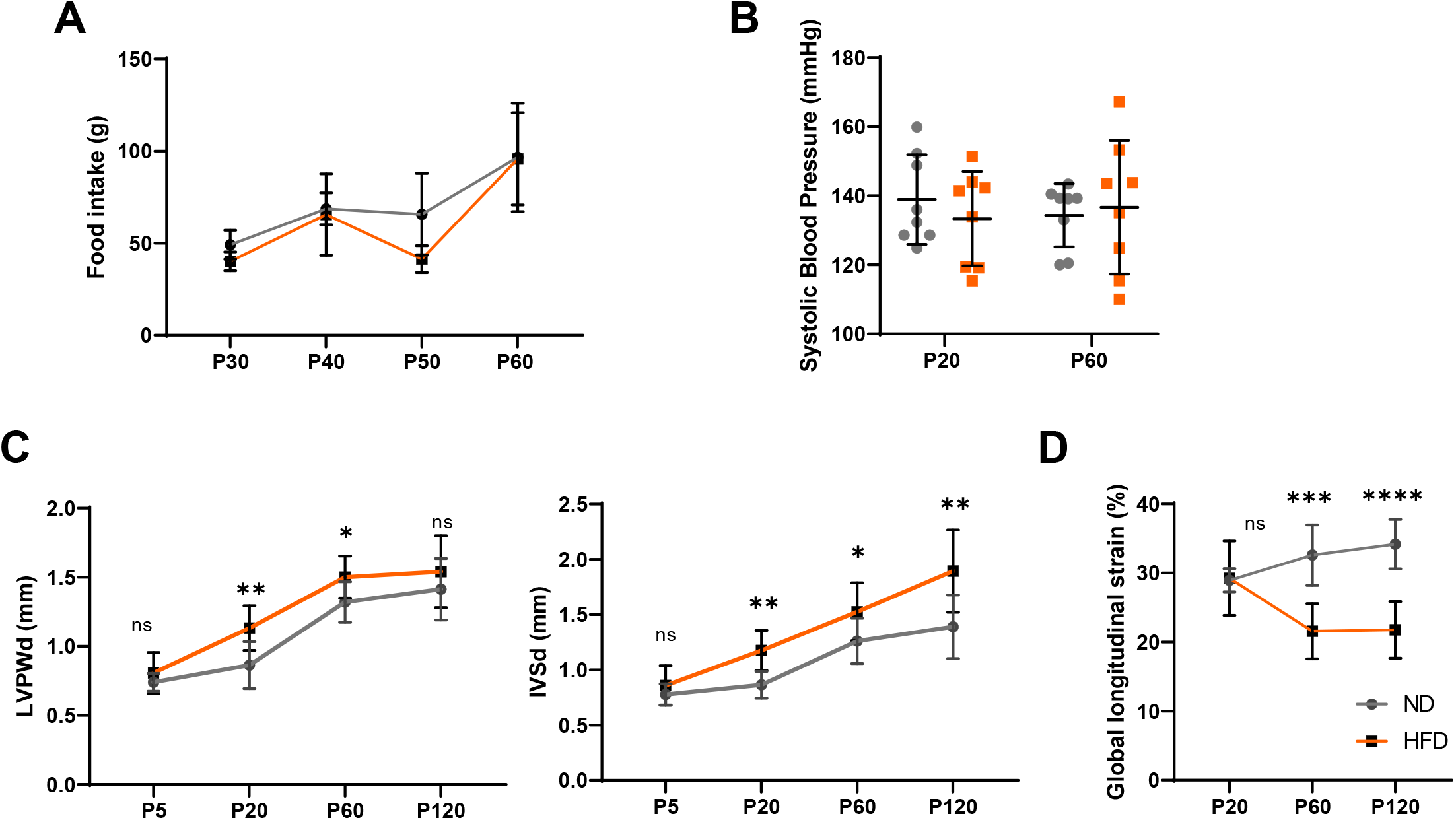
Maternal HFD induces progressive cardiac remodeling independently of postnatal food intake. **(A)** Food intake measured in normal diet (ND-Mat, grey) and high-fat diet (HFD-Mat, orange) offspring over a 2-months period (N=6 rats/group). **(B)** Systolic blood pressure measured by tail-cuff plethysmography at P20 and P60 (N=8 rats/group). **(C)** Echocardiographic assessment of cardiac morphology and function at P5, P20, P60, and P120, including left ventricular posterior wall thickness at end-diastole (LVPWd), interventricular septal thickness at end-diastole (IVSd), and **(D)** global longitudinal strain (GLS) (N=8 rats/group). Data are presented as mean±SD; two-way ANOVA followed by Tukey’s multiple comparisons test (A, B, C, D); \**P*<0.05; \*\**P*<0.01; \*\*\**P*<0.001; \*\*\*\**P*<0.0001; ns, not significant.

**Figure 2:**
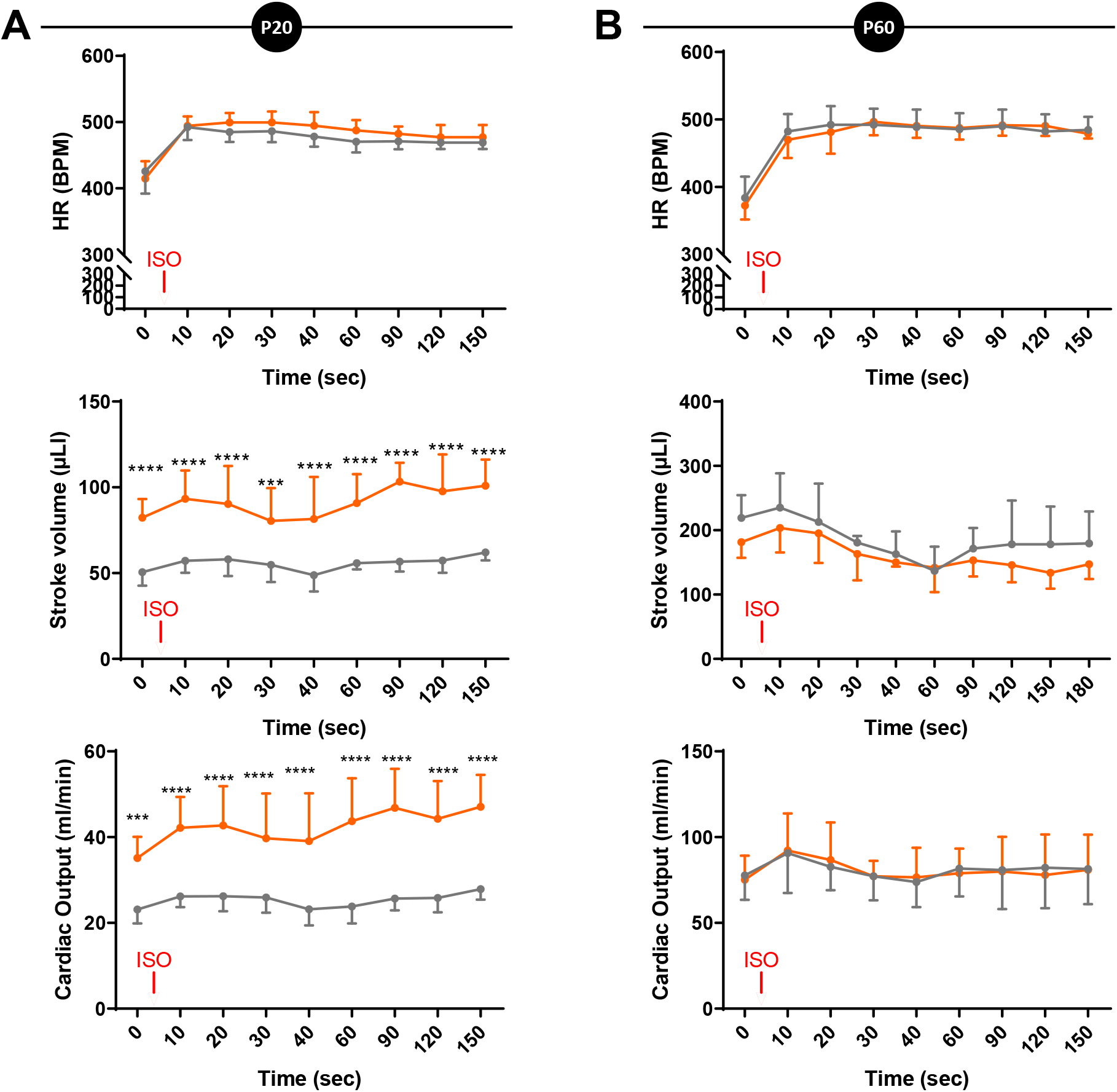
Maternal HFD-Mat enhances cardiac β-adrenergic responsiveness at P20 but not at P60. Cardiac response to acute β-adrenergic stimulation with isoproterenol, assessed by echocardiography in ND-Mat and HFD-Mat offspring at P20 **(A)** and P60 **(B)**. Heart rate (HR), stroke volume (SV) and cardiac output (CO) were measured before and after isoproterenol administration (N=8 rats/group). Data are presented as mean±SD; two-way ANOVA followed by Tukey’s multiple comparisons test; \**P*<0.05; \*\**P*<0.01; \*\*\**P*<0.001; \*\*\*\**P*<0.0001; ns, not significant.

**Figure 3:**
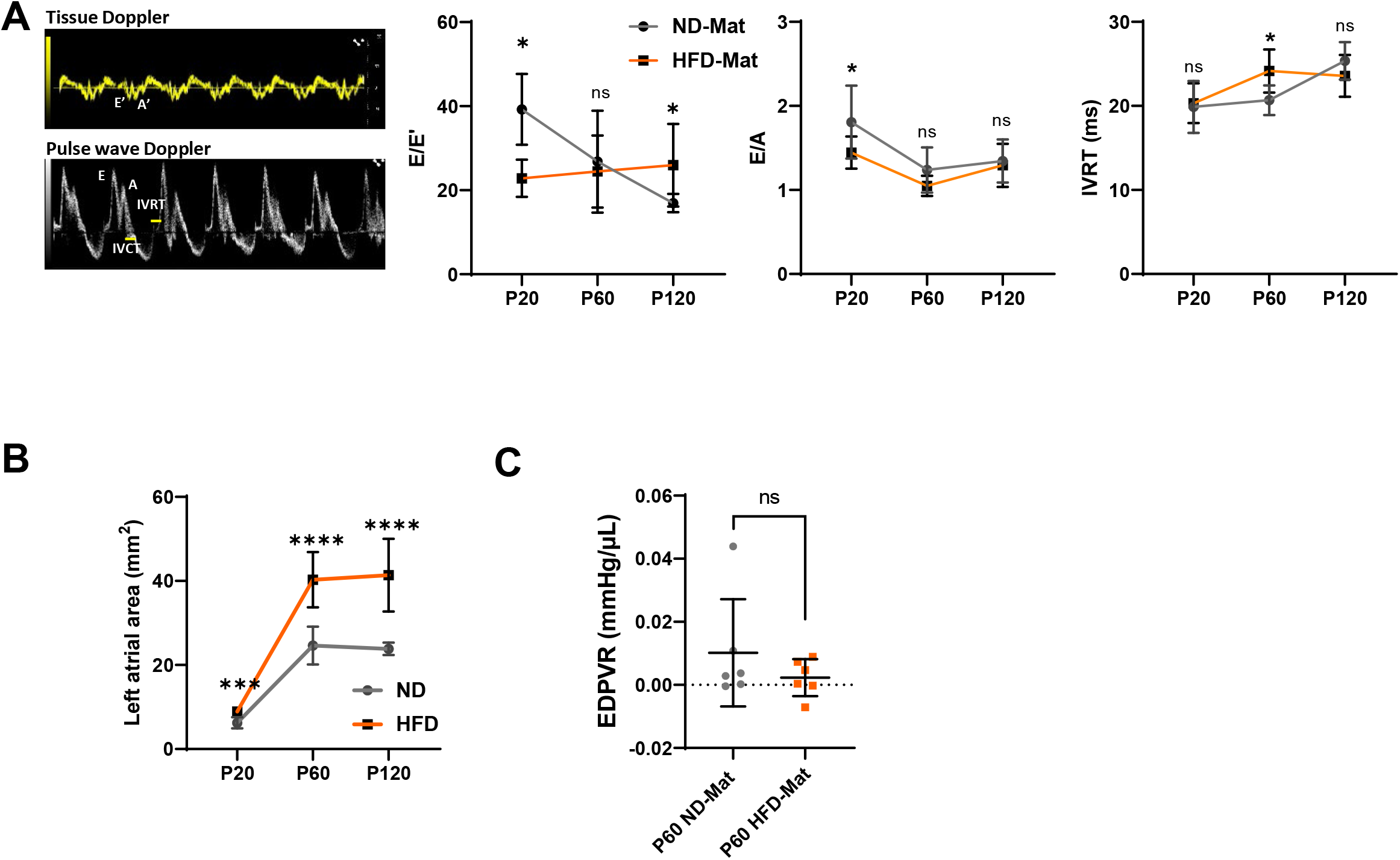
Diastolic function is largely preserved in adult HFD-Mat offspring. **(A)** Echocardiographic assessment of diastolic function by pulsed-wave Doppler and tissue Doppler imaging. Representative Doppler recordings and quantification of the E/E′ ratio, E/A ratio, and isovolumic relaxation time (IVRT) in ND-Mat and HFD-Mat offspring (N=8 rats/group). **(B)** Echocardiographic assessment of left atrial area in ND-Mat and HFD-Mat offspring at P20, P60, and P120. **(C)** Quantification of the end-diastolic pressure-volume relationship (EDPVR) obtained from pressure-volume (PV) loop analysis to evaluate passive left ventricular stiffness (N=6 rats/group). Data are presented as mean±SD; two-way ANOVA followed by Tukey’s multiple comparisons test (A, B) and Unpaired Student’s *t*-test (C); \**P*<0.05; \*\**P*<0.01; \*\*\**P*<0.001; \*\*\*\**P*<0.0001; ns, not significant.

**Figure 4.**
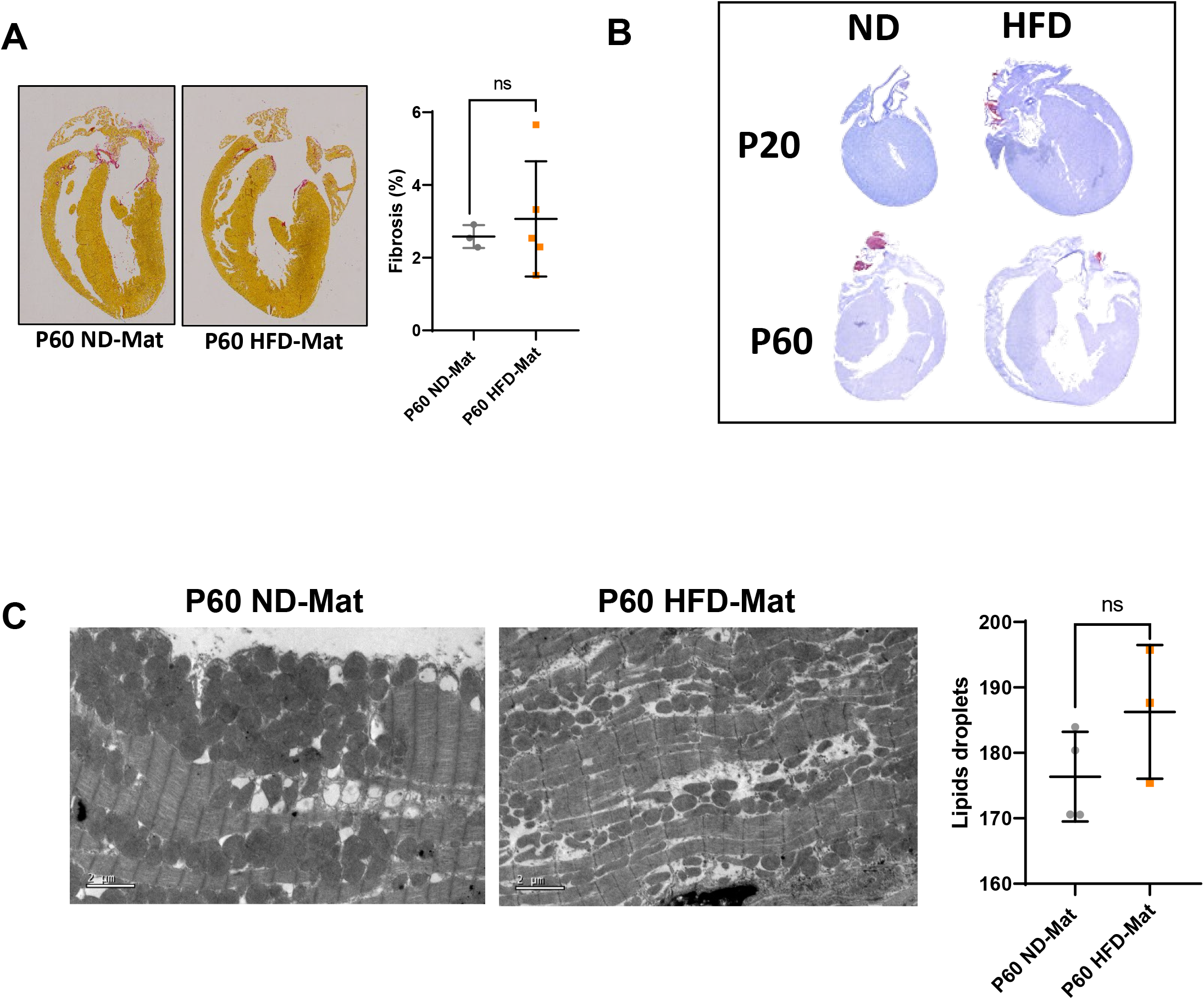
Cardiac fibrosis and lipid content are unaffected by maternal HFD in adult offspring. **(A)** Representative Picrosirius Red-stained cardiac tissue sections from ND-Mat and HFD-Mat offspring at P60, and automated quantification of fibrosis (expressed in percentage) using HistoLab software (N=3 to 4 rats/group). **(B)** Representative Oil Red O-stained cardiac tissue sections from ND-Mat and HFD-Mat offspring at P20 and P60. **(C)** Representative transmission electron microscopy images acquired at ×2,500 magnification and quantification of intracellular lipid droplets in cardiomyocytes at P60 (N=3 to 4 rats/group). Data are presented as mean±SD; unpaired *t*-test; ns, not significant.

**Figure 5:**
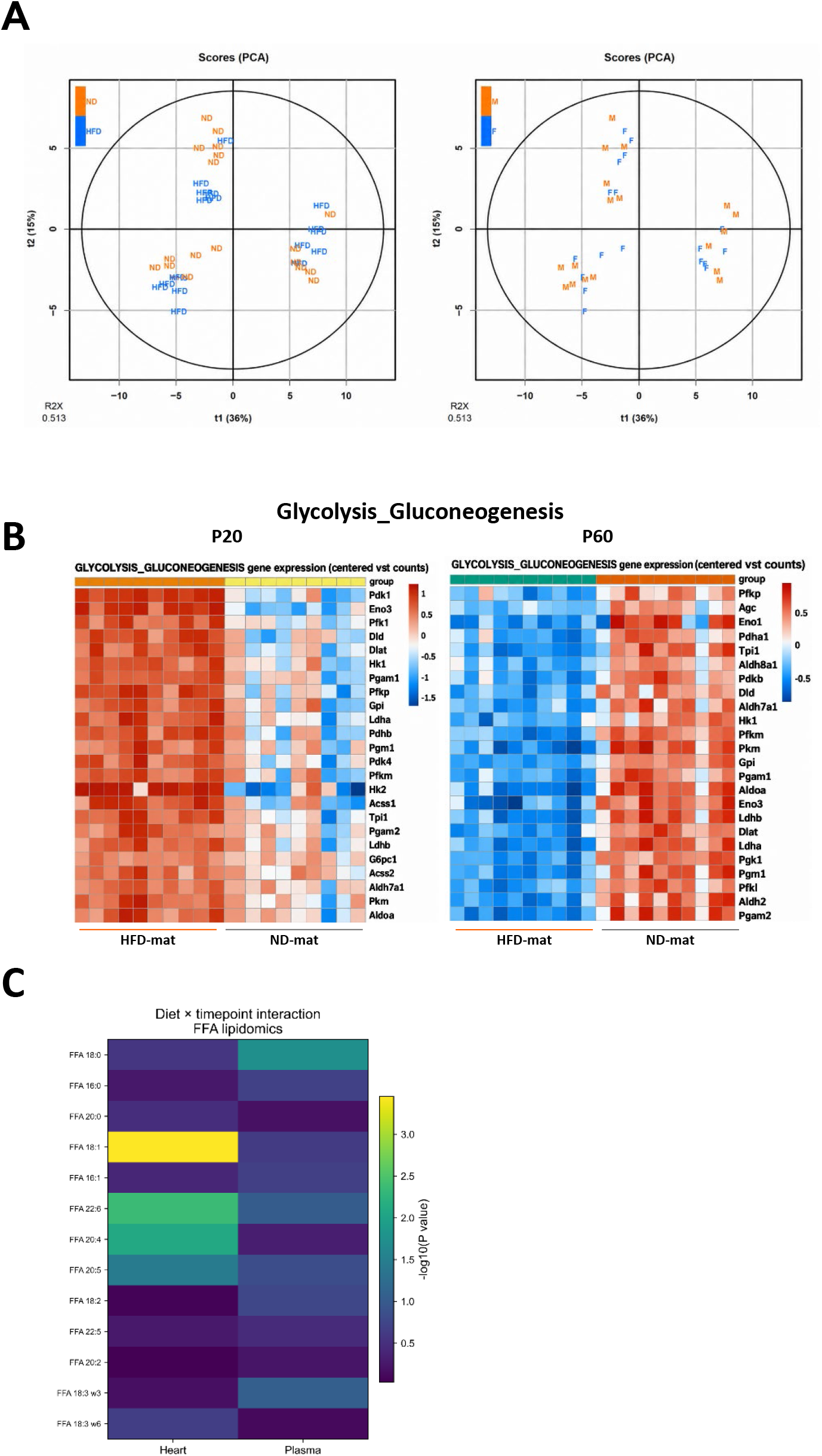
Maternal HFD reshapes the cardiac lipidome during postnatal development. **(A)** Principal component analysis (PCA) score plots of the cardiac metabolomic profiles, colored by maternal diet (ND-Mat, orange; HFD-Mat, blue; left) and by offspring sex (male, orange; female, blue; right), showing separation according to diet but not sex (N=6 rats/group). **(B)** Heatmap showing the differential abundance (log₂ fold change) of representative free fatty acids (FFAs) identified by lipidomic analysis in heart and plasma samples, highlighting metabolites significantly regulated by the diet × timepoint interaction (N=6 rats/group).

**Figure 6:**
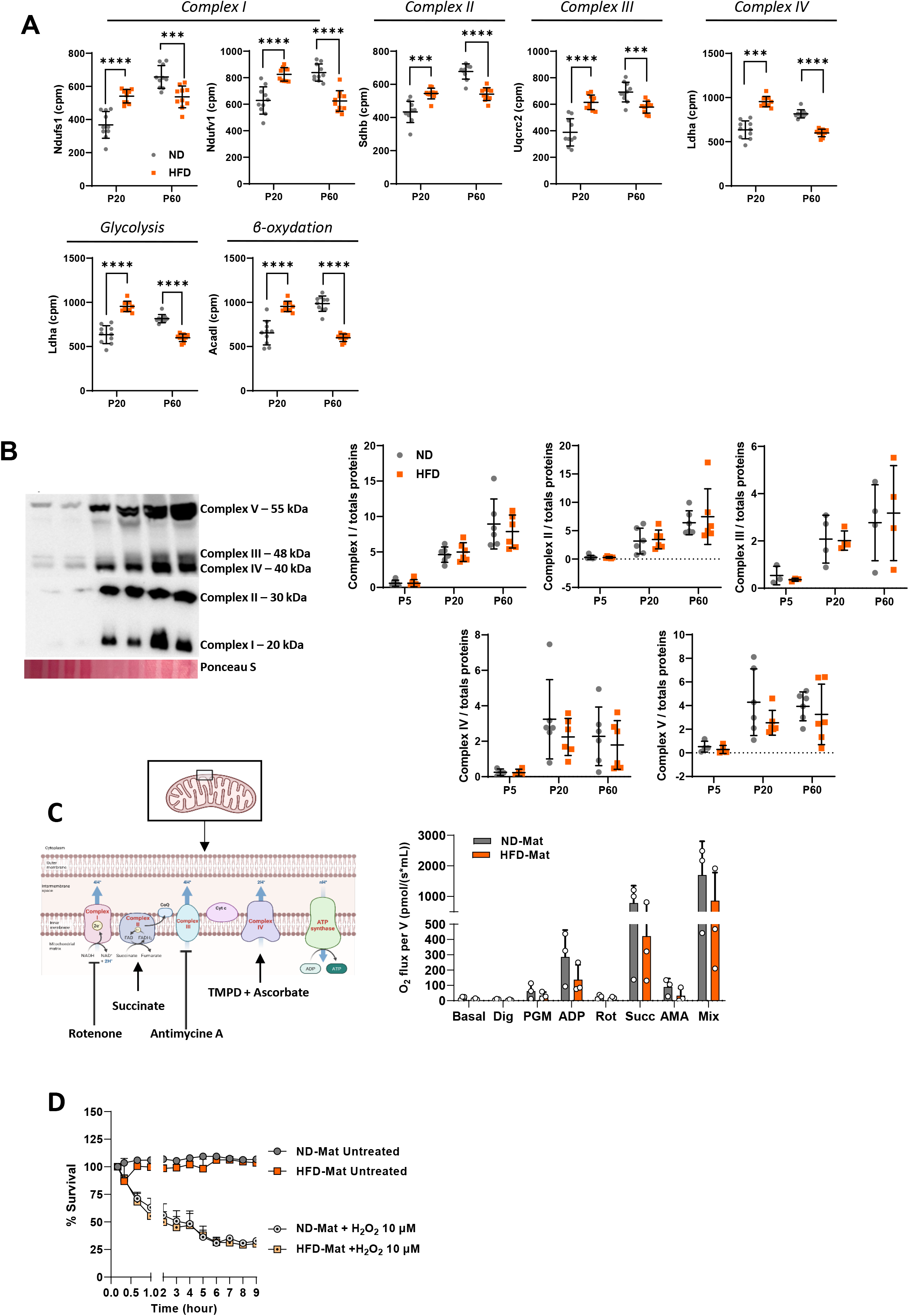
Mitochondrial oxidative metabolism is preserved in the HFD-Mat group. **(A)** RNA sequencing analysis of genes involved in mitochondrial energy metabolism in ND-Mat and HFD-Mat offspring. Expression levels, reported as counts per million (CPM), of selected genes encoding components of the mitochondrial respiratory chain (*Ndufs1*, *Ndufv1*, *Sdhb*, and *Uqcrc2*), as well as genes involved in glycolysis (*Ldha*) and fatty acid β-oxidation (*Acadl*) (N=10 rats/group). **(B)** Representative immunoblots and corresponding quantification of oxidative phosphorylation (OXPHOS) complex subunits in cardiac tissue collected from ND-Mat and HFD-Mat offspring, at P5, P20 and P60 stages (N=6 rats/group). **(C)** Mitochondrial respiratory function assessed in P60 Langendorff-isolated adult cardiomyocytes from ND-Mat and HFD-Mat offspring using high-resolution respirometry (Oroboros O2k) (N=4 rats/group). Oxygen consumption rates were measured following sequential substrate and inhibitor additions to evaluate oxidative phosphorylation capacity and electron transport chain function. **(D)** Cell survival assay following H_2_O_2_ treatment in isolated cardiomyocytes from ND-Mat and HFD-Mat offspring at P60 (N=4 rats/group). Data are presented as mean±SD; two-way ANOVA followed by Tukey’s multiple comparisons test (A, B, C) and three-way ANOVA followed by Šídák’s multiple-comparisons test; \**P*<0.05; \*\**P*<0.01; \*\*\**P*<0.001; \*\*\*\**P*<0.0001; ns, not significant.

**Figure 7.**
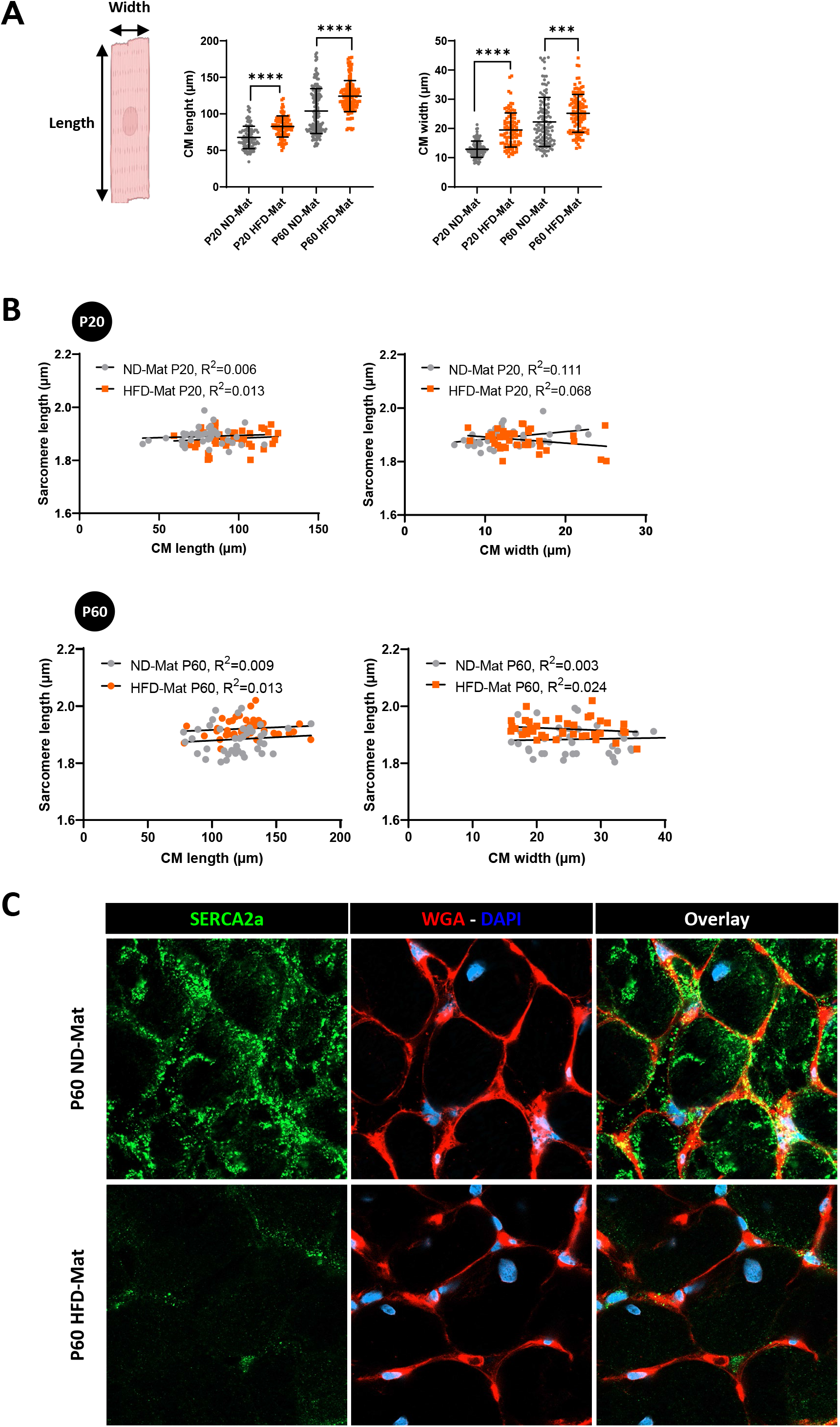
Cardiomyocyte growth and sarcomere remodeling are associated with altered SERCA2a expression. **(A)** Cell length and width measurements in Langendorff-isolated cardiomyocytes from ND-Mat and HFD-Mat offspring at P20 and P60 (N=4 rats/group; n=92 to 168 cells/group). **(B)** Correlation analyses between sarcomere length and cardiomyocyte length or width in isolated cardiomyocytes from ND-Mat and HFD-Mat offspring at P20 and P60 (N=3 rats/group, Pearson’s correlation analysis). **(C)** Representative immunofluorescence images showing SERCA2a (green) staining in fresh-frozen cardiac biopsies, transverse sections, from ND-Mat and HFD-Mat offspring at P60 (N=3 rats/group). Data are presented as mean±SD; two-way ANOVA followed by Tukey’s multiple comparisons test (A) and Pearson’s correlation analysis (B); \**P*<0.05; \*\**P*<0.01; \*\*\**P*<0.001; \*\*\*\**P*<0.0001; ns, not significant.

**Figure 8.**
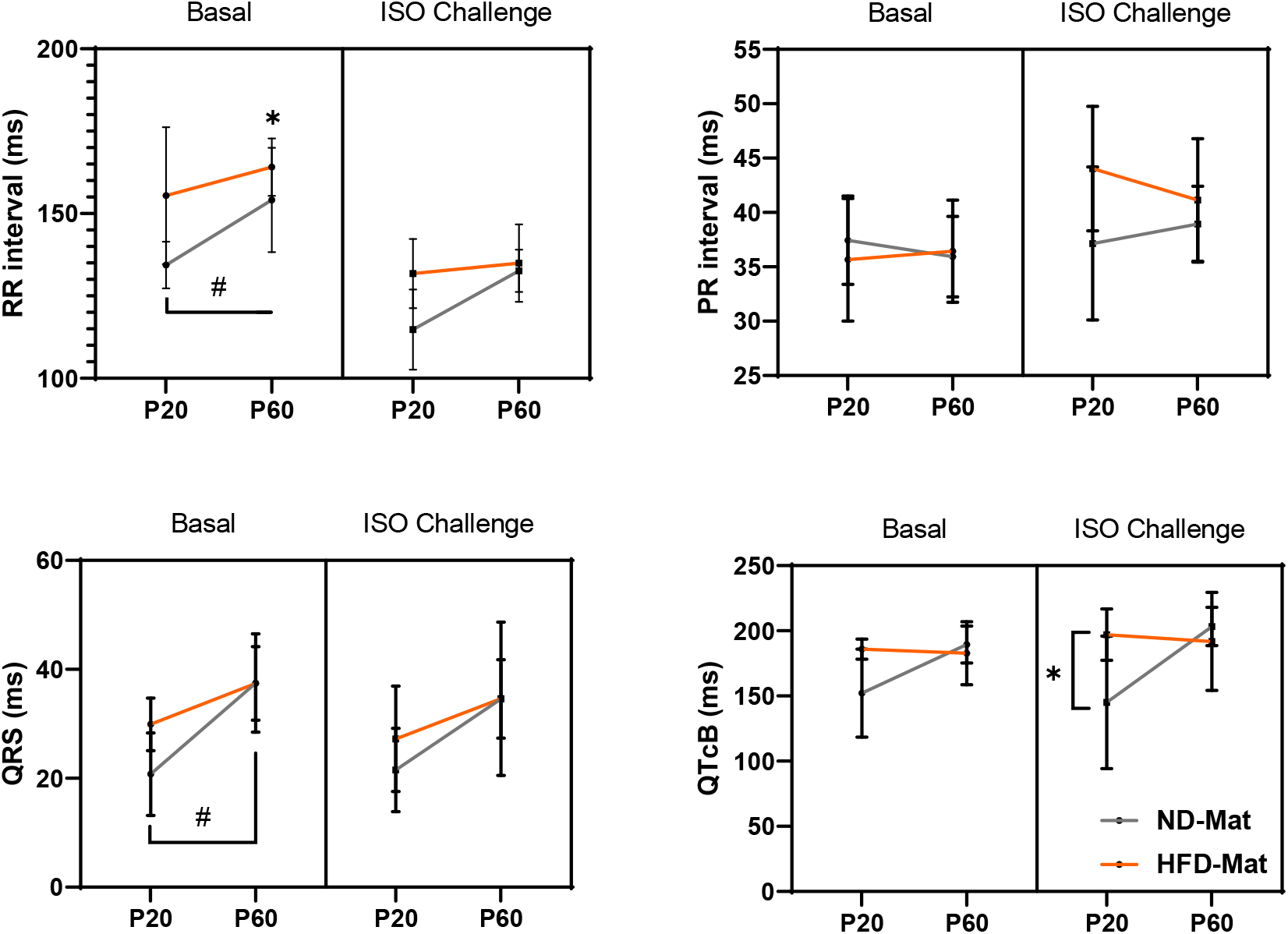
Cardiac electrical conduction and β-adrenergic reserve are preserved in HFD-Mat offspring. Electrocardiographic analysis in ND-Mat and HFD-Mat offspring at P20 and P60, under basal conditions and following isoproterenol (ISO) challenge. **(A)** RR interval. **(B)** PR interval. **(C)** QRS duration. **(D)** Heart rate-corrected QT interval, calculated using Bazett’s formula (QTcB) (N= 6 to 8 rats/group). Data are presented as mean±SD; three-way ANOVA followed by Tukey’s multiple-comparisons test; \**P*<0.05, difference within a group (developmental effect); #*P*<0.05, difference between groups (treatment effect).

## Notes

### Competing Interest Statement

The authors have declared no competing interest.

